# Near-chromosome level genome assembly of devil firefish, *Pterois miles*

**DOI:** 10.1101/2023.01.10.523469

**Authors:** Christos V. Kitsoulis, Vasileios Papadogiannis, Jon B. Kristoffersen, Elisavet Kaitetzidou, Aspasia Sterioti, Costas S. Tsigenopoulos, Tereza Manousaki

**Affiliations:** Faculty of Medicine, University of Crete, Heraklion, Greece; Institute of Marine Biology, Biotechnology and Aquaculture, Hellenic Centre for Marine Research, Heraklion, Greece; Cretaquarium, Hellenic Centre for Marine Research, Heraklion, Greece

## Abstract

Devil firefish (*Pterois miles*), a member of Scorpaenidae family, is one of the most successful marine non-native species, dominating around the world, that was rapidly spread into the Mediterranean Sea, through the Suez Canal, originating from the Indian Ocean. Even though lionfishes (Scorpaenidae) are identified among the most prosperous marine invaders, within this taxonomic group, the genomic resources are scant, while reference genome assemblies are totally absent. Here, we built and analyzed the first reference genome assembly of *P. miles* and explored its evolutionary background. The resulting genome assembly consisted of 660 contigs and scaffolds (N50 = 14,5 Mb) with a total size of about 902 Mb, while delivering 98% BUSCO completeness. We identified and described the large amount of transposable elements present in the genome and based on genomic data we constructed the first teleost phylogeny which includes a member of genus *Pterois*. The high-quality and contiguity *de novo* genome assembly built herein provides a valuable resource for future studies in species’ biology and ecology, lionfish phylogeny, the influence of transposable elements on the evolution of vertebrate genomes and fish toxins evolution.

## Introduction

The devil firefish, *Pterois miles* (Bennett, 1828), is a venomous species of the Scorpaenidae family native to the Western Indo-Pacific region, from South Africa to Red Sea and East to Sumatra (Schultz, 1986). The first occurrence of *P. miles*, as a single specimen, in the Mediterranean Sea was recorded off the Levantine coast in 1991 (Golani and Sonin, 1992), while the second, of two individuals, was almost twenty years later in the same area (Bariche et al., 2013). Soon after, the frequency of appearances, along the eastern Mediterranean Sea, rapidly increased (Crocetta et al., 2015; Kletou et al., 2016; Bilge et al., 2017; Mabruk and Rizgalla, 2019; Katsanevakis et al., 2020; Vavasis et al., 2020). Although the origin of species colonization in the Mediterranean Sea followed the invasion pattern of other Lessepsian migrants introduced from the Red Sea, through Suez Canal (Bariche et al., 2013; Dailianis et al., 2016; Kletou et al., 2016; Bariche et al., 2017; Chiesa et al., 2019; Dimitriou et al., 2019), the contribution of long-distance dispersal via aquarium trading remains a possibility (Bariche et al., 2017; Dimitriou et al., 2019). Lionfishes (genus: *Pterois*) are considered among the most thriving invaders in the history of marine invasions (Albins and Hixon, 2008) because of their rapid expansion worldwide (Azzurro et al., 2017). Indeed, the introduction in the western Atlantic of *P. miles* and of its con-generic *P. volitans*, together referred as the invasive lionfish complex (Lyons et al., 2019), is one of the fastest and most dominant marine fish introductions to date (Kletou et al., 2016, and references therein). For the Mediterranean Sea, the Suez Canal is the major pathway responsible for the spread of most of the non-indigenous species that constantly reshape its biodiversity and fishery resources (Kleitou et al., 2022). Invasive non-indigenous marine species, in general, are considered to have a major impact on local biodiversity while threatening marine industries and frequently human health (Bax et al., 2003; Arim et al., 2005; Blakeslee et al., 2019). Furthermore, they are commonly studied in evolutionary biology as models or “natural experiments” in order to explore invasion dynamics and adaptations to new niches (Barrett, 2015). Data derived from Whole Genome Sequencing (WGS) could provide promising opportunities in the exploration of potential adaptations that shape the fitness of invaders, as well as the dynamics of colonization. Further, it will provide a basis for understanding the hybrid origin of the invasive lionfishes (*P. miles* and *P. volitans*) in the Western Atlantic (Wilcox et al., 2018).

Biological characteristics of *P. miles* such as rapid somatic growth, signature anti-predatory defenses (Côté and Smith, 2018), reproductive success, discernible predatory behavior, low parasitism and ecological flexibility might explain its rapid distribution in the Mediterranean Sea (Dailianis et al., 2016), whereas the species population is rising rapidly along the coastlines (Kletou et al., 2016). Yet, few genomic resources of the species are available, including the mitochondrial genome (Dray et al., 2016) and DNA barcoding data (Chiesa et al., 2019, and references therein).

Devil firefish belongs to Scorpaenidae, a large family of venomous marine species including lionfishes, scorpionfishes and stonefishes (Diaz, 2015). Their venom (toxins) is mainly secreted from spines that are present in dorsal, lateral, pelvic and anal fins. These toxins are composed by two subunits α and β to form their active dimeric structure (Kiriake and Shiomi, 2011; Kiriake et al., 2013; Chuang and Shiao, 2014; Campos et al., 2021). The excreted venom is used for defensive purposes alongside other strategies (Campos et al., 2021, and references therein) that lead to successful anti-predatory adaptations. These scorpionfish toxins have multiple biological activities and their range differs between different species, despite their high similarity and conservation in specific domains (Chuang and Shiao, 2014; Campos et al., 2021). So far, toxins from stonefishes (stonustoxin, verrucotoxin and neoverrucotoxin) have been mainly identified and characterized (Ghadessy et al., 1996; Ueda et al., 2006; Kiriake and Shiomi, 2011; Kiriake et al., 2013), while toxins from other genera (*Scorpaena*, *Scorpaenopsis*, *Inimus* and *Pterois*) were recognized by similarity and cloning using the previous ones (Kiriake and Shiomi, 2011; Kiriake et al., 2013; Chuang and Shiao, 2014; Xie et al., 2019; Campos et al., 2021). However, the scorpionfish toxins are relatively understudied, even though there is a high diversity between them (Xie et al., 2019). Due to the absence of genomic data inside this family, the origin and evolution of toxins within scorpaenid species, and specifically in genus *Pterois*, still remains ambiguous.

The aim of this study was to construct, annotate and analyze the first high-quality genome assembly of *P. miles*. Through a combination of Oxford Nanopore Technologies (ONT), Pacific Biosciences (PacBio) and Illumina reads, we explored the genomic background of such a successful and unique invader, the devil firefish. Being the first top-quality genome sequenced within the Scorpaenidae family, this valuable resource could provide a critical conveyance to unveil and highlight the insights of species’ biology, ecology and phylogeny in further investigations of invasive traits across the Mediterranean Sea.

## Materials & Methods

### Sample collection, DNA and RNA library construction and sequencing

Animal care and handling were carried out following well established guidelines (Degrazia and Beauchamp, 2019).

A specimen was caught alive in Bali (North coast), Crete, Greece and was transferred in the indoor facilities of Cretaquarium (part of the Hellenic Centre for Marine Research, HCMR). The specimen was kept alive for three weeks in a 150 L tank with semi-closed circuit with airlift, mechanical and biological filtration, water recirculation rates of 30 and 100%/h. Temperature was adjusted at 19 °C +/- 1,0 °C and the photoperiod was set to 12hL:12hD. The specimen was anesthetized using clove oil, and blood was collected with a sterilized and heparinized syringe from the caudal vein of the fish. Blood was kept in sterilized and heparinized 1.5 ml tubes on ice, until DNA extraction.

DNA was extracted, for the purpose of ONT sequencing, from 4ul of blood with the Qiagen Genomic Tip 100/G kit. DNA integrity was assessed by electrophoresis in 0.4 % w/v megabase agarose gel. Three ligation libraries (SQK-LSK109) were constructed using 2.3 microgram DNA as input for each library. The libraries were sequenced on two R9.4.1 flow cells, on the in-house MinION Mk1B and MinION Mk1C sequencers, respectively. The resulting fast5-files were basecalled with Guppy v5.0.11, using the “sup” (super-accuracy) configuration and a minimum quality score of 7.

For the Illumina sequencing, the same procedure and protocol were used for DNA extraction. The template DNA for Illumina sequencing was sheared by ultrasonication in a Covaris instrument. A PCR-free library was prepared with the Kapa Hyper Prep DNA kit with TruSeq Unique Dual Indexing.

For transcriptomic data, total RNA was extracted from seven tissues (brain, gonad, gills, heart, liver, muscle and spleen). Tissue grinding in liquid nitrogen with pestle and mortar was followed by the sample homogenization and lysis in TRIzol buffer (Invitrogen) according to the manufacturer’s guidelines. The quantity and the quality of the extracted RNA was estimated with NanoDrop ND-1000 spectrophotometer and with agarose gel (1.5%) electrophoresis, as well as with Agilent 2100 Bioanalyzer using RNA 6000 pico kit. The library preparation was conducted using Pacific Biosciences protocol for Iso-Seq™ Express Template Preparation for Sequel and Sequel II Systems. The seven libraries (one for each tissue) were sequenced on a PacBio sequel II instrument, on one SMRT cell.

Illumina and Pacific Biosciences library preparation and sequencing were carried out by the Norwegian Sequencing Centre (www.sequencing.uio.no), hosted in the University of Oslo.

### Genomic data pre-processing

The length filtering and adapter trimming of basecalled ONT reads were carried out with Porechop v0.2.4 (https://github.com/rrwick/Porechop) using default parameters, adding the extra parameter “–discard_middle” to prune reads with potential inner adapters. The quality control was performed using Nanoplot v.1.20 (De Coster et al., 2018). The quality assessment of Illumina reads was performed with FastQC v0.11.9 (Andrews, 2010) while both filtering of low quality reads and adapter trimming, using Trimmomatic v0.39 (Bolger et al., 2014). The reads were processed by Trimmomatic on a 4-base sliding window with an average cutting-off threshold score lower than 15 Phred score. Leading and trailing bases with a quality score less than 10 Phred were trimmed out, while reads shorter than 75 bp and average score lower than 30 Phred were removed.

### *De novo* genome assembly

For the *de novo* genome assembly, using a hybrid approach, the long reads from ONT were combined with short and highly accurate Illumina reads. At first, the ONT reads were used for the construction of the initial draft *de novo* assembly and the first rounds of polishing, while the Illumina reads were used for the later rounds of polishing. The draft assembly was built from ONT reads using the *de novo* assembler Flye v2.9. (Kolmogorov et al., 2019), which uses a repeat graph as core data structure, with default parameters and a genome size estimation of 900Mb. The genome size estimation was based on the corresponding entry of *Pterois volitans* on www.genomesize.com. Then, the draft assembly was polished in two rounds with RACON v1.4.3 (Vaser et al., 2017) using the filtered long reads, mapped against it by Minimap v2.22 (Li, 2018) which resulted in the intermediate assembly. Further polishing was performed on the intermediate genome assembly by Medaka v1.4.4 (https://github.com/nanoporetech/medaka) and then with Pilon v1.23 (Walker et al., 2014) after mapping with Minimap v2.22 (Li, 2018) the preprocessed Illumina reads against the assembly taken from Medaka, resulting in the final reference genome assembly used for all downstream analyses.

The draft, intermediate and final assemblies were evaluated by two commonly used criteria: (i) the N50 statistic from contig sizes, using QUAST v.5.0.2 (Gurevich et al., 2013), and (ii) the completeness score based on the presence of universal single copy ortholog genes, using BUSCO v.5.3 (Manni et al., 2021) against Actinopterygii ortholog dataset 10 (actinopterygii_odb10). BUSCO was run with default parameters adding the extra parameter “--augustus” to enable species-specific training for gene prediction by AUGUSTUS v.3.4 (Stanke et al., 2008). Alternative values (e.g. L90) were calculated (Table 1) with a custom python tool, ELDAR (https://github.com/ckitsoulis/ELDAR).

**Table 1.**
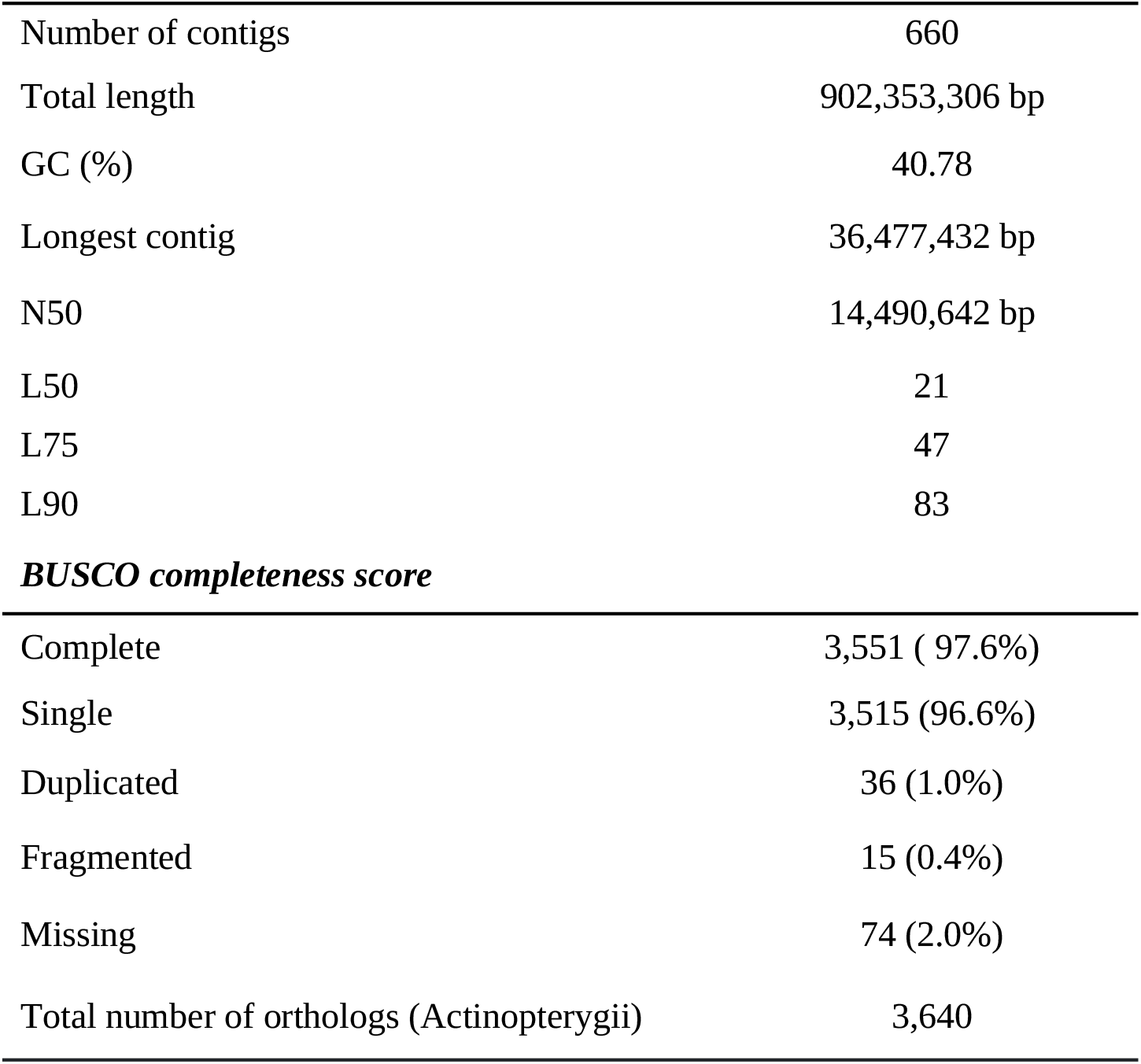
Polished genome assembly statistics and completeness.

The whole genome assembly pipeline which was used in the present study was previously designed by Danis et al. (2021), containerized by Angelova et al. (2022) (https://github.com/genomenerds/SnakeCube) and ran in the IMBBC High performance computing (HPC) facility “Zorbas” (Zafeiropoulos et al., 2021).

### Genome annotation

#### Transposable elements annotation

A *de novo* transposable elements (TEs) library was generated from the constructed genome assembly of *P. miles*, using the Extensive *de novo* TE Annotator (EDTA) package (Ou et al., 2019), an automated whole-genome TE annotation pipeline, with default parameters. In our case, the RepeatModeler2 (Flynn et al., 2020) was utilized to additionally support the identification of TE families inside the EDTA algorithm, using the extra parameter “-- sensitive 1”. The non-redundant TE library was then separated into three sub-libraries based on its TEs classification, so far, using a custom python script “library_split.py”: i) Classified TE sequences, ii) Unclassified TE sequences in the level of superfamily (partially classified), and iii) Unclassified - Unknown TE sequences. The two latter sub-libraries were classified again using DeepTE (Yan et al., 2020), a transposon classification tool which depends on convolutional neural networks (CNN). The annotation probability threshold was strictly set to 0.8 (“-prop_thr 0.8”). A step of header correction and reformation, using bash commands, in every sub-library occurred before their concatenation to the final TE annotated library, in order to achieve a compatible format for the next steps. Finally, RepeatMasker v4.1.2 (Tarailo-Graovac and Chen, 2009) performed the initial TE annotation and genome soft- masking, utilizing the NCBI/RMblast search engine, based on the previously-described library. To achieve a more accurate and detailed annotation/categorisation of TE (Table 2), based on an up-to-date TE classification system (Makalowski et al., 2019), a python-based parser “RM_parser.py” was developed and used for the output files of RepeatMasker. The designed workflow is presented schematically in Supplementary Figure 1.

**Table 2.**
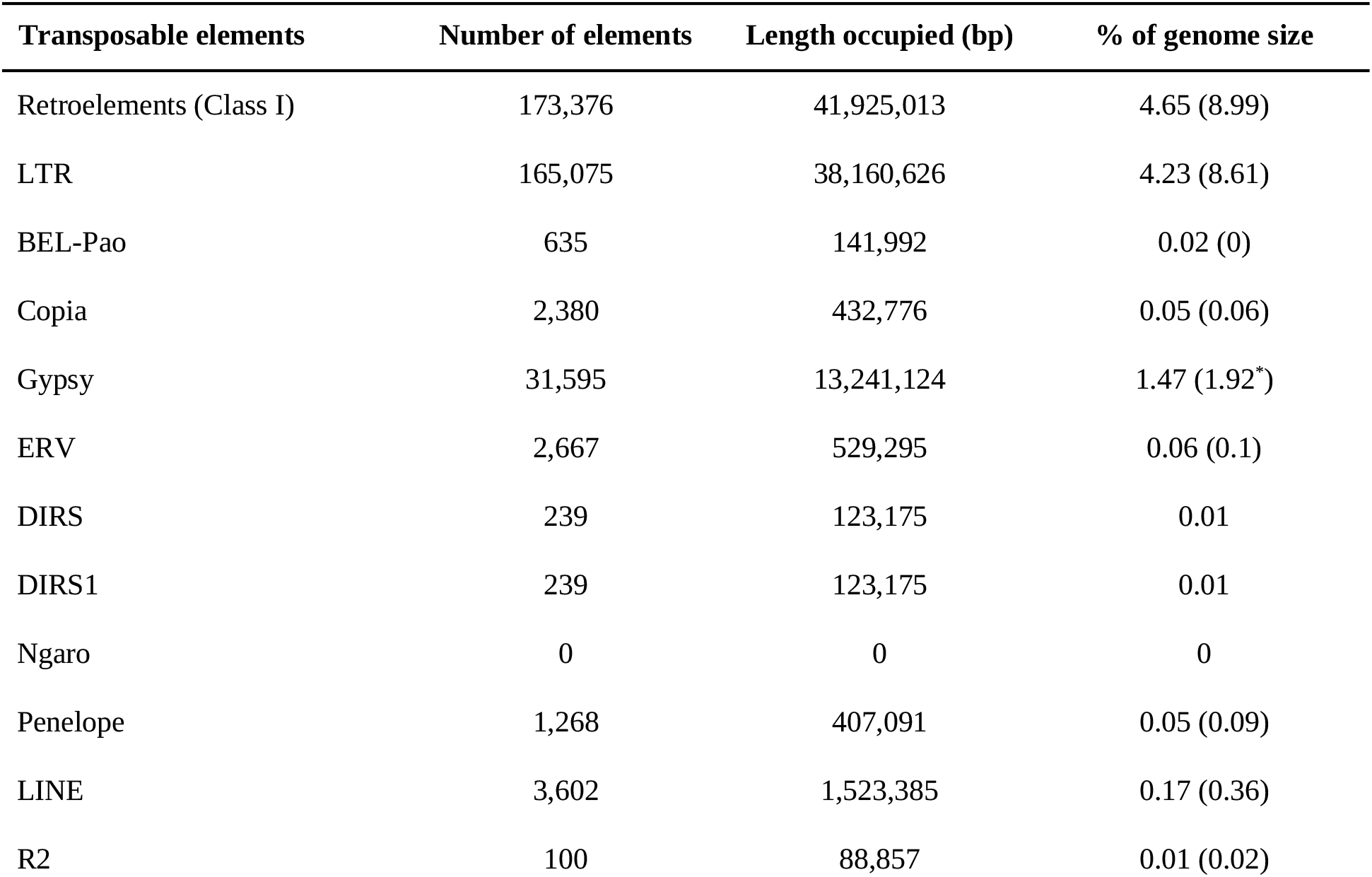
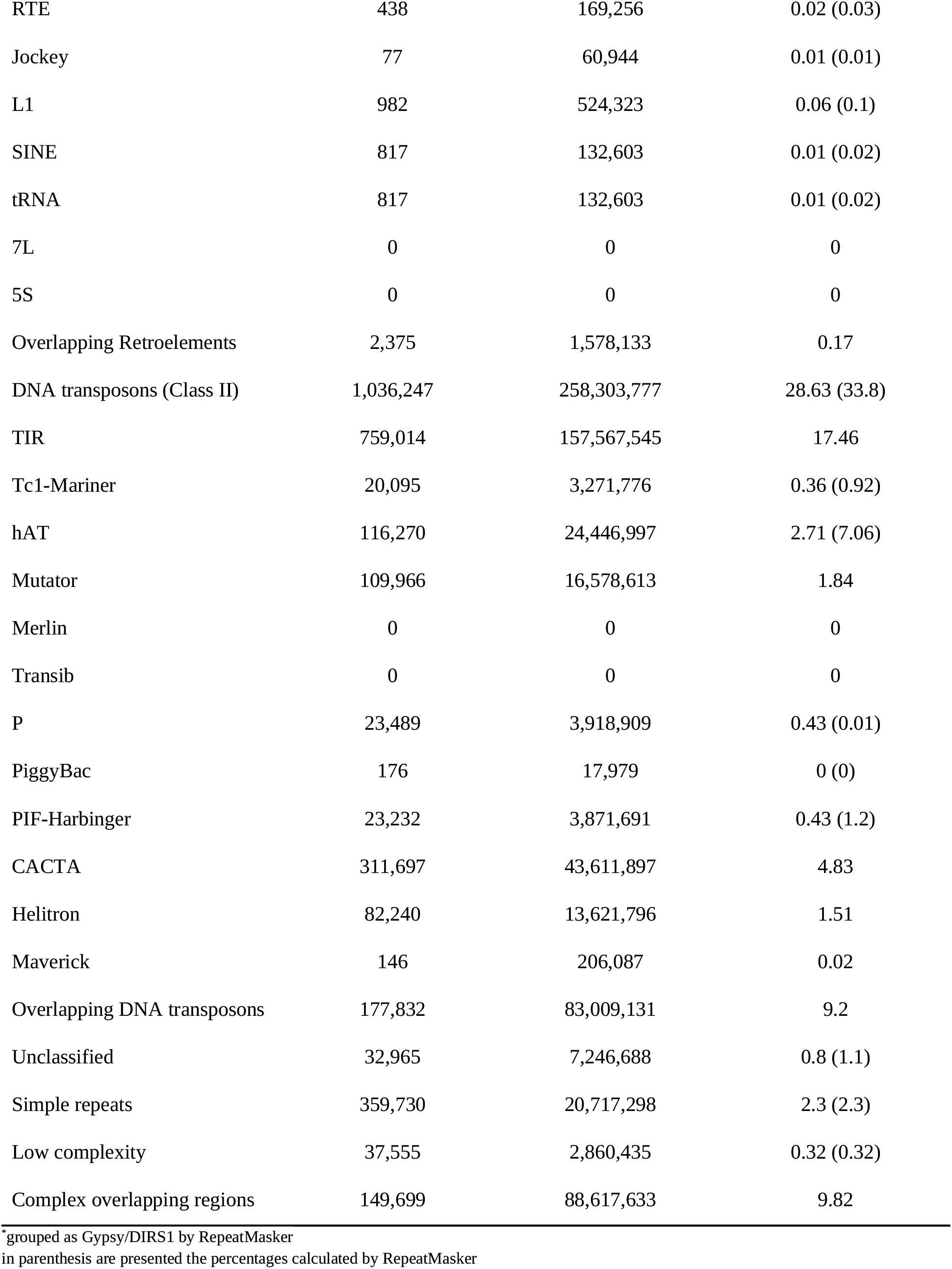
Transposable element annotation statistics.

| Transposable elements | Number of elements | Length occupied (bp) | % of genome size |
| --- | --- | --- | --- |
| Retroelements (Class I) | 173,376 | 41,925,013 | 4.65 (8.99) |
| LTR | 165,075 | 38,160,626 | 4.23 (8.61) |
| BEL-Pao | 635 | 141,992 | 0.02 (0) |
| Copia | 2,380 | 432,776 | 0.05 (0.06) |
| Gypsy | 31,595 | 13,241,124 | 1.47 (1.92*) |
| ERV | 2,667 | 529,295 | 0.06 (0.1) |
| DIRS | 239 | 123,175 | 0.01 |
| DIRS1 | 239 | 123,175 | 0.01 |
| Ngaro | 0 | 0 | 0 |
| Penelope | 1,268 | 407,091 | 0.05 (0.09) |
| LINE | 3,602 | 1,523,385 | 0.17 (0.36) |
| R2 | 100 | 88,857 | 0.01 (0.02) |
| RTE | 438 | 169,256 | 0.02 (0.03) |
| Jockey | 77 | 60,944 | 0.01 (0.01) |
| L1 | 982 | 524,323 | 0.06 (0.1) |
| SINE | 817 | 132,603 | 0.01 (0.02) |
| tRNA | 817 | 132,603 | 0.01 (0.02) |
| 7L | 0 | 0 | 0 |
| 5S | 0 | 0 | 0 |
| Overlapping Retroelements | 2,375 | 1,578,133 | 0.17 |
| DNA transposons (Class II) | 1,036,247 | 258,303,777 | 28.63 (33.8) |
| TIR | 759,014 | 157,567,545 | 17.46 |
| Tc1-Mariner | 20,095 | 3,271,776 | 0.36 (0.92) |
| hAT | 116,270 | 24,446,997 | 2.71 (7.06) |
| Mutator | 109,966 | 16,578,613 | 1.84 |
| Merlin | 0 | 0 | 0 |
| Transib | 0 | 0 | 0 |
| P | 23,489 | 3,918,909 | 0.43 (0.01) |
| PiggyBac | 176 | 17,979 | 0 (0) |
| PIF-Harbinger | 23,232 | 3,871,691 | 0.43 (1.2) |
| CACTA | 311,697 | 43,611,897 | 4.83 |
| Helitron | 82,240 | 13,621,796 | 1.51 |
| Maverick | 146 | 206,087 | 0.02 |
| Overlapping DNA transposons | 177,832 | 83,009,131 | 9.2 |
| Unclassified | 32,965 | 7,246,688 | 0.8 (1.1) |
| Simple repeats | 359,730 | 20,717,298 | 2.3 (2.3) |
| Low complexity | 37,555 | 2,860,435 | 0.32 (0.32) |
| Complex overlapping regions | 149,699 | 88,617,633 | 9.82 |

\*grouped as Gypsy/DIRS1 by RepeatMasker in parenthesis are presented the percentages calculated by RepeatMasker

#### Structural annotation - gene prediction

For transcriptome data, circular consensus sequencing (CCS) reads were generated using the CCS application on SMRT Link v10.2 and then Iso-Seq analysis was performed on them, with default parameters, using the Iso-Seq pipeline, IsoSeq v3.4 (https://github.com/PacificBiosciences/IsoSeq), until the production of high quality (HQ) consensus full-length transcripts. The IsoSeq pipeline included three basic steps: i) generation of CCS reads, ii) classification of full-length (FL) reads, and iii) clustering of full-length non- contatamer (FLNC) reads to obtain high-quality consensus transcripts. The number of resulting intermediate reads to final isoforms in the IsoSeq pipeline are presented in Table 3. The HQ transcripts were aligned (spliced-wise) against the soft-masked genome assembly of *P. miles*, using GMAP v2021.08.25 (Wu and Watanabe, 2005). SAM files were sorted and converted to BAM using samtools v1.15.1 (Danecek et al., 2021), and the redundant transcripts were finally collapsed to generate a non-redundant HQ full-length transcripts set, using cDNA Cupcake v28.0 (https://github.com/Magdoll/cDNA_Cupcake) with a minimum alignment coverage equal to 0.99 and a minimum alignment identity of 0.95.

**Table 3.** Number of reads from CCS to high-quality isoforms.

| Tissue | CCS | HiFi | FLNC (polyA) | HQ isoforms |
| --- | --- | --- | --- | --- |
| muscle | 682,522 | 651,026 | 679,844 | 34,862 |
| liver | 863,535 | 814,531 | 861,396 | 50,651 |
| heart | 747,994 | 704,226 | 743,228 | 73,273 |
| brain 1 | 16,449 | 15,739 | 15,986 | 2,467 |
| brain 2 | 33,032 | 32,232 | 32,810 | 7,042 |
| gonad | 396,510 | 377,750 | 395,080 | 17,804 |
| spleen | 596,858 | 568,143 | 594,852 | 66,103 |
| gills | 193,697 | 184,914 | 192,470 | 25,831 |
| consensus isoforms |  |  |  | 124,307 |

Gene prediction was conducted based on a hybrid strategy of transcriptome-based (non-redundant high quality transcripts), homology-based (curated protein sets) and *ab initio* methods, using a semi-automated workflow consisting of 12 individual tools and intermediate custom python and bash scripts (Supplementary Figure 2). In the first step, HQ isoforms and curated proteomes of 20 actinopterygian species were aligned (spliced-wise) to the soft-masked genome assembly. The non-redundant HQ trascriptome set was previously aligned using GMAP v2021.08.25 (Wu and Watanabe, 2005). For protein homology evidence, a BLAST database was generated from protein sequences of 20 species (Table 4), being downloaded from UniProtKB/Swiss-Prot (https://www.uniprot.org/), using DIAMOND v2.0.14 (Buchfink et al., 2015). In the second step, Mikado v2.3.3 (Venturini et al., 2018) was used, a python-based pipeline which identifies the “best” set of transcripts from multiple sources, in order to return potential gene models from the transcriptome and protein homology evidence. Homology evidence for each of the predicted transcripts provided to Mikado were generated based on the BLAST DB, using DIAMOND v2.0.14 (Buchfink et al., 2015) while open reading frame (ORF) predictions of Mikado-selected transcripts were produced by Transdecoder v5.5 (https://github.com/TransDecoder/TransDecoder). All information and evidence were merged afterwards to generate the most accurate evidence-based gene models, using Mikado steps “serialise” and “pick”. These gene models had been used in later steps of gene prediction and annotation update. In the third step, Augustus v3.4 (Stanke et al., 2008) was trained with two optimization rounds on a subset of gene models (generated in step 2) that fulfilled specific criteria: i) full length, ii) non-redundant, iii) over a blast hit score of 0.5, and iv) with at least 2 exons. The training set was selected using a custom python script “select_training.py”. To take advantage of Augustus ability to incorporate hints (gene, protein, intron etc) for generating high confident gene models, species-specific exon hints and spliced protein alignments were generated and merged, secondarily. For exon hints, the exons coordinates were extracted from the previously produced annotation file using python scripts. For the spliced protein alignments, three well annotated protein sets of species *Oryzias latipes* (downloaded from UniProtKB), *Gasterosteus aculeatus* (downloaded from Ensembl) and *Argyrosomous regius* (Papadogiannis et al., 2023) were aligned to the genome assembly, using Exonerate v2.4 (https://github.com/nathanweeks/exonerate). The annotation files were merged, sorted and then filtered for exonic evidence extraction using python. *Ab initio* prediction on the *P. miles* genome assembly, alongside the generated hints, was performed by Augustus v3.4 (Stanke et al., 2008) with extra parameters “--allow_hinted_splicesites=atac” and “--alternatives-from-evidence=false”. In the fourth step, gene models, generated in the previous steps (from Mikado and Augustus), were merged into a consensus gene set, after two updating rounds, using PASA v2.4.1 (Haas et al., 2003), an eukaryotic genome annotation pipeline. For this reason, Mikado-predicted protein coding gene models were loaded into PASA to create the initial MySQL DB of transcripts, the Augustus predictions were loaded to the DB and it was updated later on with the Mikado-predicted genes. The same procedure was followed, as a second updating round, starting this time from the resulting annotation of the first round. In the last step, genes were filtered to remove predictions with in-frame STOP codons and those that overlapped with TEs. For the first case, the gene models were cleaned for potential identical isoforms using Agat (https://github.com/NBISweden/AGAT), the artifacts were recognised using gffread (https://github.com/gpertea/gffread), while they were removed with bash commands. For the second case, candidate models were found using bedtools v2.30 (Quinlan and Hall, 2010) “intersection” command, with a minimum overlapping score of 0.5 “-f 0.50” and filtered out with bash commands as well. The completeness evaluation of transcripts and genes, in each step, was performed using BUSCO v5.3 (Manni et al., 2021) against the Actinopterygii ortholog dataset 10 (Table 5).

**Table 4.** Species included in protein homology BLAST database.

| Scientific name | Common name | UniProt ID | Number of proteins | Reference |
| --- | --- | --- | --- | --- |
| *Amphilophus citrinellus* | Midas cichlid | UP000261340 | 31,742 | Yunyun et al. (2022) |
| *Amphiprion ocellaris* | Clown anemonefish | UP000257160 | 31,745 | Ryu et al. (2022) |
| *Astyanax mexicanus* | Blind cave fish | UP000018467 | 39,383 | McGaugh et al. (2014) |
| *Betta splendens* | Siamese fighting fish | UP000515150 | 41,617 | Fan et al. (2018) |
| *Carassius auratus* | Goldfish | UP000515129 | 82,968 | Chen et al. (2019) |
| *Clupea harengus* | Atlantic herring | UP000515152 | 37,255 | Kongsstovu et al. (2019) |
| *Danio rerio* | Zebrafish | UP000000437 | 46,841 | Howe et al. (2013) |
| *Esox lucius* | Northern pike | UP000265140 | 71,519 | Rondeau et al. (2014) |
| *Gymnodraco acuticeps* | Ploughfish | UP000515161 | 39,915 | Bista et al. (2022) |
| *Haplochromis burtoni* | Burton's mouthbrooder | UP000264840 | 34,332 | Brawand et al. (2014) |
| *Hippocampus comes* | Tiger tail seahorse | UP000264820 | 27,735 | Lin et al. (2016) |
| *Ictalurus punctatus* | Channel catfish | UP000221080 | 40,203 | Wang et al. (2022) |
| *Lepisosteus oculatus* | Spotted gar | UP000018468 | 22,463 | Braasch et al. (2016) |
| *Oreochromis niloticus* | Nile tilapia | UP000005207 | 74,622 | Conte et al. (2017) |
| *Oryzias latipes* | Japanese rice fish | UP000001038 | 36,128 | Kasahara et al. (2007) |
| *Perca flavescens* | American yellow perch | UP000295070 | 21,644 | Feron et al. (2020) |
| *Salmo salar* | Atlantic salmon | UP000087266 | 82,390 | Lien et al. (2016) |
| *Sparus aurata* | Gilthead sea bream | UP000472265 | 69,200 | Perez-Sanchez et al. (2019) |
| *Takifugu rubripes* | Japanese pufferfish | UP000005226 | 51,078 | Kai et al. (2011) |
| *Xiphophorus maculatus* | Southern platyfish | UP000002852 | 35,279 | Schartl et al. (2013) |

**Table 5.** Basic statistics of predicted gene models.

| Type | Number | Mean size (bp) | Longest (bp) | Shortest (bp) | Genome size (%) |
| --- | --- | --- | --- | --- | --- |
| gene | 24,645 | 15,518 | 445,933 | 201 | 42.37 |
| transcript | 25,040 | 15,440 | 445,933 | 201 | 42.84 |
| 5' UTR | 13,119 | 228 | 10,216 | 1 | 0.24 |
| exon | 231,331 | 207 | 14,732 | 3 | 5.33 |
| CDS | 227,256 | 159 | 6,853 | 16 | 4 |
| intron | 206,291 | 1,609 | 188,322 | 21 | 36.05 |
| 3' UTR | 11,114 | 914 | 14,608 | 5 | 1.07 |

#### Functional annotation

The functional annotation of *P. miles* predicted gene set was performed using three different strategies and tools. The first approach was based on similarity search (reciprocal hits) against the annotated genes of zebrafish (*D. rerio*) using BLASTp v2.12+ (Altschul et al., 1990) with parameters: “-evalue 1e-6”, “-max_target_seqs 1” and “-hdps 1”, in order to reduce the number of queries for the later annotation update. In the second one, results were fetched with EggNOG-mapper v2.1.7 (Cantalapiedra et al., 2021) based on fast orthology assignments using pre-computed clusters and phylogenies from eggNOG v5.0 database (Huerta-Cepas et al., 2019). For the last approach, annotations were retrieved using PANNZER2 (Törönen and Holm, 2022), a weighted k-nearest neighbour classifier which is based on SANSparallel (Koskinen and Holm, 2012) for homology similarity against UniProt and enrichment statistics, using various user-defined scoring functions. Prediction of gene names, Gene Ontology (GO) annotations, KEGG pathway IDs, Pfam domains and descriptions from all aforementioned strategies were filtered and assigned to gene models using a custom python script “FUNfilter.py”. Gene names, in each case, were selected based on the most frequent occurrence, while KEGG pathway IDs and Pfam domains were obtained directly from EggNOG-mapper. An additional step was performed for GO terms (biological process) being mapped to gene models, by using the assigned gene names as queries and retrieving terms from UniProtKB (https://www.uniprot.org/) with “Retrieve/ID mapping tool”, for human, mouse and zebrafish. Finally, GO terms resulted as a set of terms between those predicted by EggNOG-mapper v2.1.7 and UniProtKB.

### Phylogenomic analyses

#### Orthology assignment

To identify orthologous and paralogous genes, 46 whole-genome protein coding gene sets (longest isoforms) from teleost species (Supplementary Table 1), along with the one of *P. miles*, were compared using OrthoFinder v2.5.2 (Emms and Kelly, 2019), with default parameters. The initial dataset of genomes was collected from Genomes-NCBI Datasets (https://www.ncbi.nlm.nih.gov/datasets/genomes/) and Ensembl (https://www.ensembl.org/) based on the following criteria: i) genomes at the chromosome and/or scaffold level of contiguity and ii) N50 > 10Mb. The longest isoform per gene was selected initially (in GFF format) using Agat (https://github.com/NBISweden/AGAT) and then extracted as FASTA file with gffread (https://github.com/gpertea/gffread). Each proteome set was assessed for completeness using BUSCO v.5.3 (Manni et al., 2021) against Actinopterygii ortholog dataset 10. For the final set, only proteomes which exceeded a predefined completeness threshold (90%) were included and only one species per genus was kept for the final analysis.

#### Species tree inference

The phylogenetic hierarchical orthogroups (HOGs) produced by OrthoFinder were filtered, at first, to select those with complete representation from *P. miles* and then, only those containing a single gene copy per species, to exclude potential paralogs. Afterwards, orthogroups which had a representation from at least 43 out of 47 species (> 91.4%) were selected, using a custom python script (aragorn_orthoX.py). The protein sequences of each HOG were aligned using MAFFT v7.505 (Katoh and Standley, 2013). In cases of HOGs with no representation from certain species, the corresponding sequences were filled with gaps equal to the total length of the HOG alignment using a custom python script (gimli_clean.py). Then, all aligned HOG sequences were concatenated into a superalignment matrix with bash commands. The initial matrix was trimmed to remove spurious sequences and poorly aligned regions using trimAl v1.4.1 (Capella-Gutiérrez et al., 2009), with default parameters and strict mode. ModelTest-NG v0.1.7 (Darriba et al., 2019) was used for the selection of the best-fit model and IQTREE v2.2.0.3 (Minh et al., 2020) for the maximum-likelihood phylogenetic tree inference. To assess the confidence of branches, IQ-TREE was run for 1000 bootstrap replicates (ultrafast bootstrap mode). The phylogenetic tree was finally visualized using R/RStudio Team (2022) and package “ggtree” (Yu, 2020) within a custom R script (tree.R), selecting *Lepisosteus oculatus* and *Polyodon spathula*, as an outgroup clade.

### Comparative genomic analysis

#### Synteny analysis

Synteny analysis was performed at the gene level between *P. miles* and three-spined stickleback, *G. aculeatus*. For this purpose, one-to-one orthologues were selected from the HOGs produced earlier by OrthoFinder, to compare the physical localization of genetic loci within species. The 42 longest contigs of *P. miles* genome assembly (representing ∼73.5% of the genome size) were selected for visualization against the 21 chromosomes of *G. aculeatus*, using Circos (Krzywinski et al., 2009).

#### Gene families expansion and contraction

Changes in gene family size (expansions and contractions) were estimated using CAFE v5 (Mendes et al., 2020). The HOGs summary table of all species from OrthoFinder was retrieved and modified earlier by a custom python script “aragorn_orthoX.py”, resulting in a count matrix of genes per species and family, in order to be used by CAFE. Following CAFE’s developers instructions, HOGs absent in more than 10 species out of 47 and the ones in which the difference between the maximum and minimum number of genes was greater than 70 (*max_i_(n_genes_) - min_i_(n_genes_) > 70*), were filtered out from the analysis. An ultrametric binary tree was produced with R package “ape” (Paradis and Schliep, 2019) in a custom R script “tree_calibration.R”. For that, we used the phylogenetic tree, produced earlier, and the divergence times taken from TIME-TREE (http://www.timetree.org/) between 4 different species’ combinations, *Polyodon spathula* - *Danio rerio*, *Danio rerio* - *Takifugu rubripes*, *Oryzias latipes* - *Mola mola* and *Dicentrarchus labrax* - *Mola mola*. CAFE was finally run using 3 different gamma function categories (-k) to estimate λ parameter (corresponding to the rate of change of families), 400 iterations (-I) and a p-value equal to 0.05 (-pvalue). After the analysis, we selected for visualization only the Perciformes clade, as a subset of the phylogenetic tree.

#### Duplication event estimation

To infer gene duplication events in *P. miles* from gene family trees and the estimated phylogenetic tree, we used GeneRax v2.0.4 (Morel et al., 2020). Initially, protein sequences from each HOG, produced by OrthoFinder, were aligned to each other using MAFFT v7.505 (Katoh and Standley, 2013) and trimmed with trimAl v1.4.1 (Capella-Gutierrez et al., 2009), in strict mode. Orthogroups with less than three sequences were excluded from the following procedure. The best-fit model was estimated from each HOG and later was used for the inference of a single maximum-likelihood gene tree, using IQTREE v2.2.0.3 (Minh et al., 2020). After manually inspecting and correcting the estimated substitution models for some cases, gene trees and their models along with the phylogenetic species tree were used to estimate duplication events with GeneRax.

#### Gene ontology terms descriptive analysis

For GO terms descriptive analysis, firstly we downloaded the core ontology (OBO format) from Gene Ontology DB (http://geneontology.org/docs/download-ontology/), and then the predicted gene set of *P. miles* and their assigned GO terms were mapped with GO biological process descriptions provided by the core ontology, using a custom python script “obo_mapper.py”. Then, these GO terms and their descriptions were grouped/mapped into the gene families of their genes (HOGs from OrthoFinder), which were previously identified as rapidly expanding from CAFE and involved in duplication events by GeneRax.

#### Toxin gene evolution in lionfishes

To identify genes responsible for the secreted toxin of devil firefish, toxins (proteins) identified in other scorpaenid species (Ghadessy et al., 1996; Ueda et al., 2006; Kiriake and Shiomi, 2011; Kiriake et al., 2013; Chuang and Shiao, 2014; Xie et al., 2019) were downloaded from NCBI (Table 6) and aligned to the genome of *P. miles*, using tBLASTx v.2.12 (Altschul et al., 1990). All proteins included were about 700 amino acids long and constituted of three exons. The identified coding regions on the genome of *P. miles* fulfilling specific criteria (a. blast hit against all proteins (toxins), b. non-overlapping to each other, c. with three potential exons) were translated into proteins using similarity results from BLAST and ExPASy Translate tool (Gasteiger, 2003), after the recognition of correct ORFs. The protein sequences were, then, aligned against with MAFFT v7.505 (Katoh and Standley, 2013) and trimmed using trimAl v1.4.1 (Capella-Gutiérrez et al., 2009), in strict mode. Finally, the alignment was manually inspected using Jalview (Waterhouse et al., 2009). ModelTest-NG v.0.1.7 (Darriba et al., 2019) was used for the estimation of the substitution model and IQTREE v2.2.0.3 (Minh et al., 2020) to infer the maximum-likelihood phylogenetic tree. The final unrooted tree was visualized with FigTree v.1.4.4 (http://tree.bio.ed.ac.uk/software/figtree/).

**Table 6.**
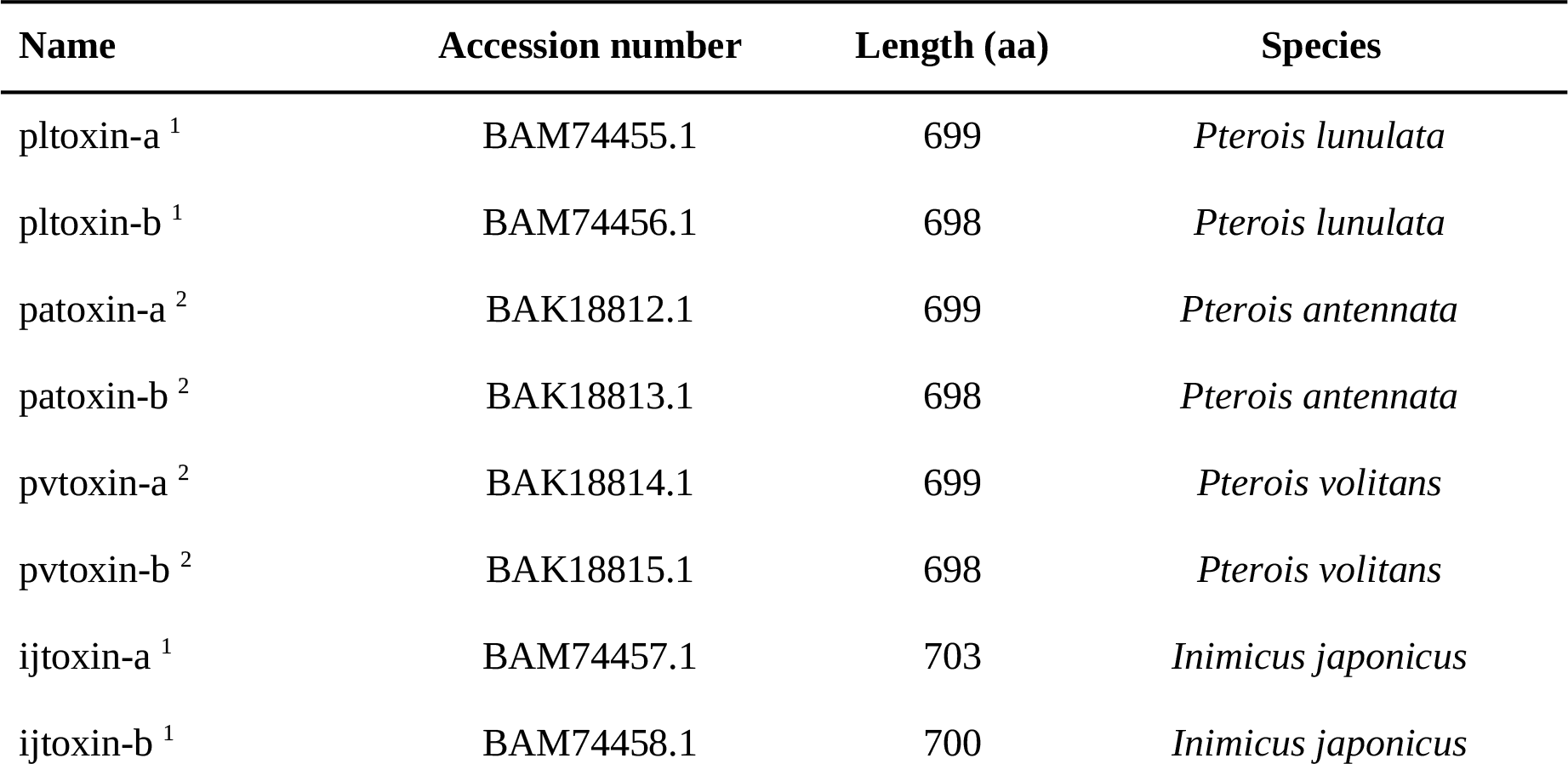
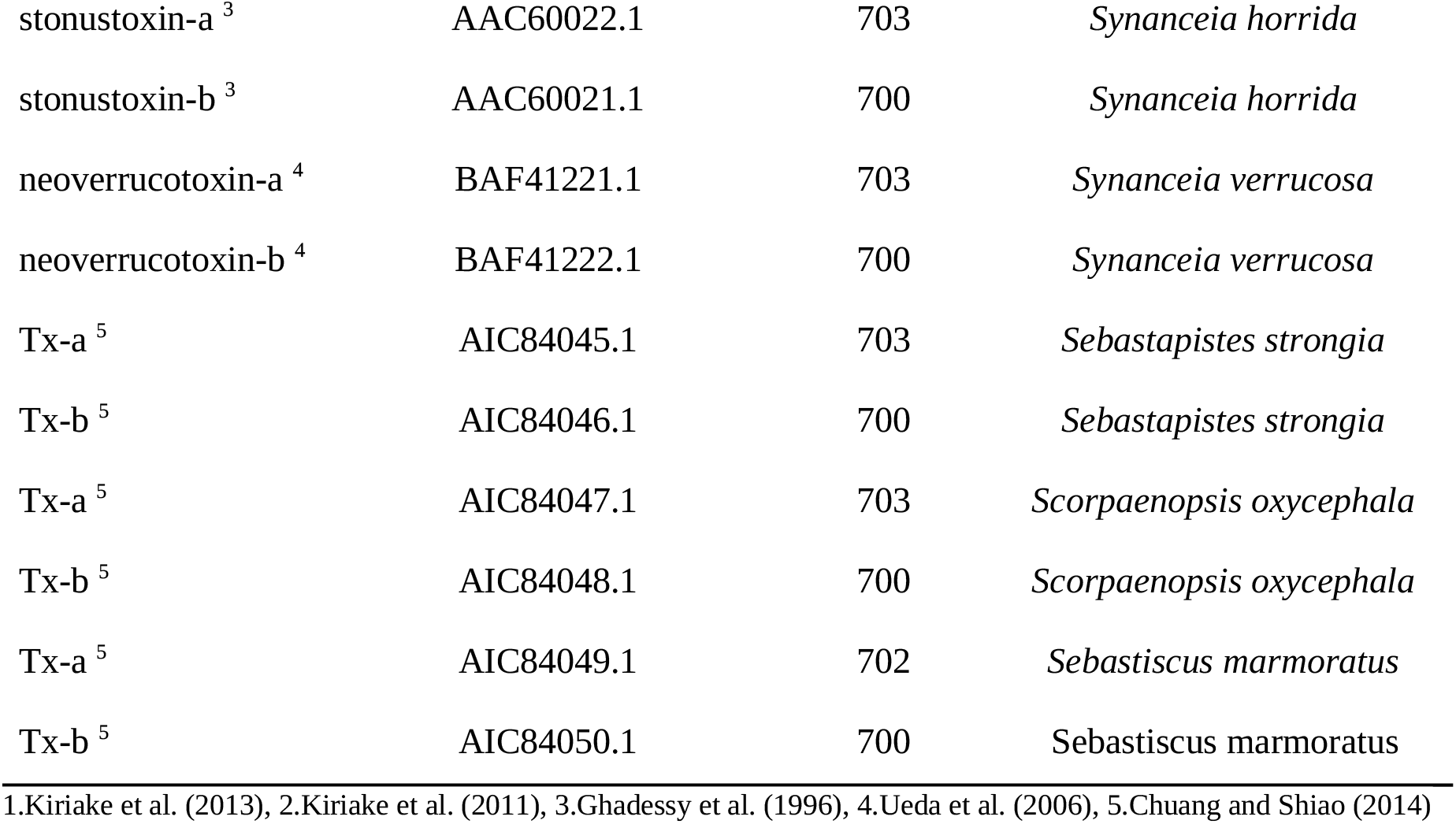
Scorpionfish toxins downloaded from NCBI.

## Results

### Genomic sequencing results

Sequencing yielded a total of 38.66 Gb of raw genomic ONT reads, from which 36.16 Gb had a Phred quality score above Q7, as well as 3.75 Gb of raw Illumina reads. After the pre- processing steps of both trimming and quality filtering, 35.72 Gb of ONT, for the initial assembly, and 3.16 Gb of Illumina reads, for later polishing, were maintained for the downstream process of *de novo* construction of the genome assembly (Table 7).

**Table 7.** Summary of genomic sequencing throughput.

| Sequencing technology | Raw reads | Quality-controlled reads | Coverage |
| --- | --- | --- | --- |
| Illumina | 24,836,074 | 20,980,358 | 3.5 x |
| MinION | 2,245,868 | 2,224,738 | 39.5 x |

### Genome size and assembly completeness

The final genome assembly contained 660 contigs with a total length of about 902.3 Mb. The longest contig was sized at 36.5 Mb and the N50 statistic value at 14.5 Mb (Table 1). At least, 90% of the genome size was represented in the 83 longest contigs of the produced assembly (Supplementary Figure 3). The GC content of the genome was calculated at 40.78% (GC-rich regions at the 42 longest contigs are presented in Supplementary Figure 4). For genome completeness assessment, 3,566 out of 3,640 BUSCO genes were present (98%), against the Actinopterigyian ortholog dataset (v.10). Of those, 3,551 genes (97.6%) were complete, while only 74 (2.0%) were missing (Table 1), suggesting a high level of contiguity and completeness of the *de novo* genome assembly.

### Genome annotation

#### Transposable elements annotation

About 46.5% of the genome assembly (∼ 416.7 Mb) in *P. miles*, consisted of transposable elements (Figure 1). Class I Retroelements make up 4.65% of the genome assembly and LTR order is the most dominant with a representation of at least 4.23%, with its superfamily Gypsy of 1.47%. Class II of TEs (DNA transposons) represents a high amount (28.6%) of the whole genome, while elements of the TIR order and its superfamily CACTA were mostly found, with 17.46% and 4.83% respectively, among the high confidence and completely classified DNA TEs. Additionally, 9.8% of the masked genome are regions of complex composition of overlapping TEs, not clearly defined as discrete elements during masking (Table 2). The distribution of TE content in the 42 longest contigs is presented in Supplementary Figure 4.

**Figure 1.**
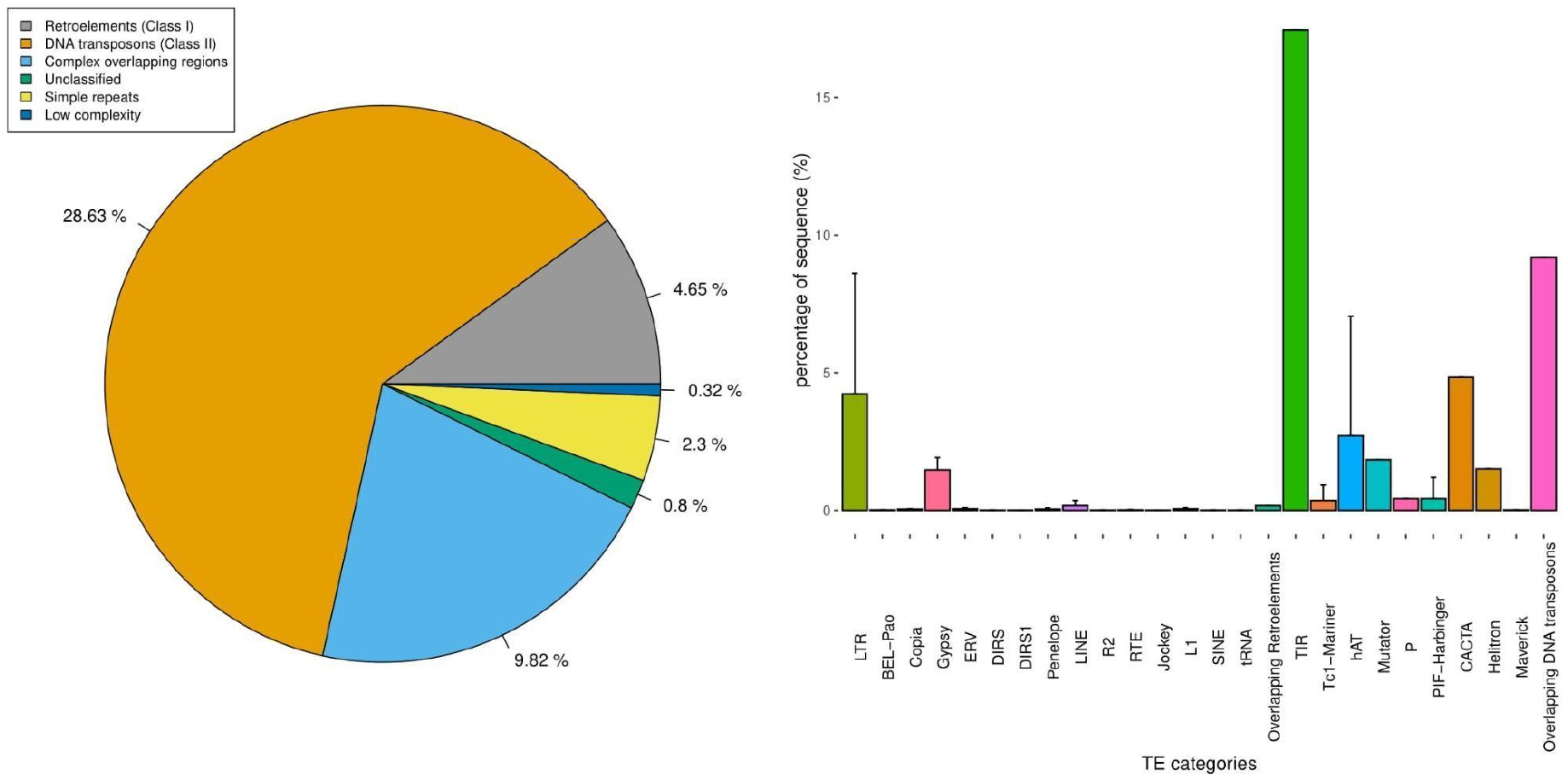
Percentage of transposable element categories representation in the genome of *P. miles*.

#### Transcriptome analysis

From 6,245,243 initial CCS reads, Iso-seq analysis yielded a total of 124,307 HQ consensus isoforms. The samples with the higher number of transcripts were the heart, spleen and liver (Table 3), and they shared almost 6,500 of them (Supplementary Figure 5). However, the total amount of HQ transcripts shared between all sampled tissues was notably low (∼1,000). From the total number of HQ transcripts, 91,666 were aligned properly to the assembled genome (representing 73.74%) and used as evidence for the gene prediction.

#### Structural and Functional annotation

The hybrid approach of HQ full-length transcripts-based, homology-based and *ab initio*- based methods resulted in a total of 25,410 candidate protein-coding gene models. We filtered out genes with in-frame stop codons (266 putative genes) and those overlapping with TEs (505 gene models). We ended up with 24,639 potential gene models (Supplementary_file1_annotation.gff), representing in size 382.3 Mb of the genome assembly (Table 5). A subset of 22,473 genes were assigned with gene names and 23,521 matched at least one functional description, accounting 89.7% and 95.4% of the total number of genes, respectively. In terms of GO, KEGG pathway IDs and PFAM domains, 22,115 (88.3%), 15,071 (61.1%) and 20,003 (81.1%) genes were annotated, in each case (Supplementary_file2_functional.tsv).

From a core set of 3,640 single-copy orthologous genes (Actinopterigyii lineage, odb 10), 3,414 (93.8%) were found to be present in the predicted gene set (Table 8), with 3,233 (88.8%) identified as complete (3,082 as single-copy and 151 as duplicated) and 181 (5.0%) as fragmented, while 224 (6.2%) of them were not present, using BUSCO v5.3 (Manni et al., 2021).

**Table 8.**
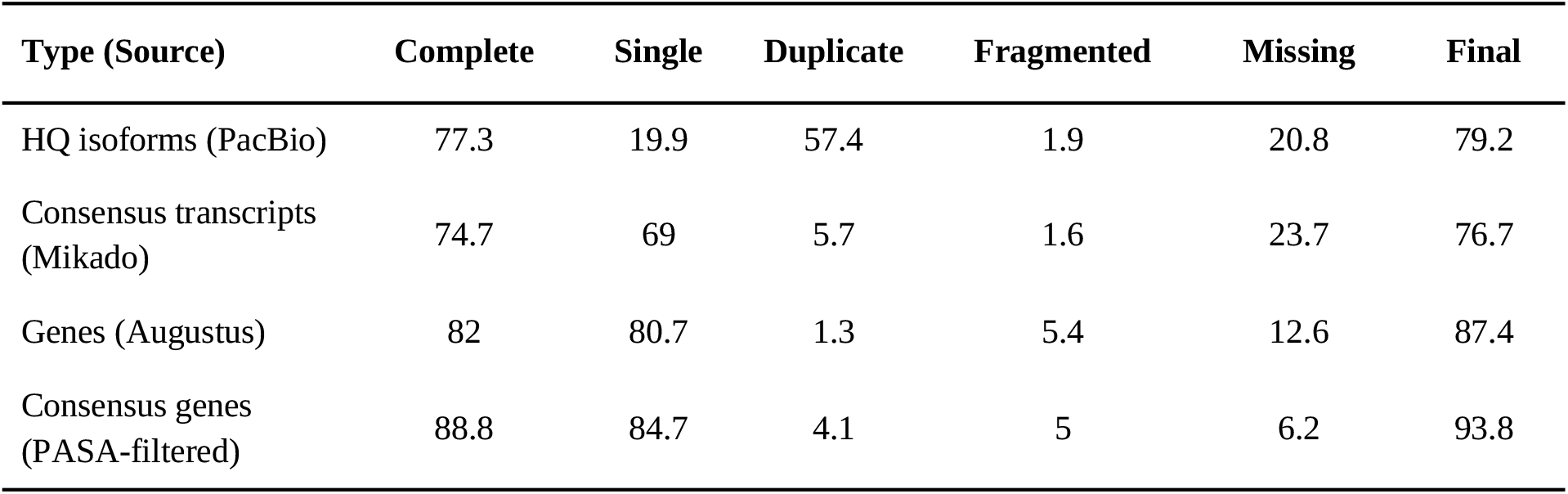
BUSCO completeness assessment (%) in each step of structural annotation.

### Orthology assignment and phylogenomic analysis

The total number of proteins included in the proteomes of all 47 teleost fish species (Supplementary Table 1) and analyzed by OrthoFinder, was 1,108,753 and 97.8% of them were assigned to 28,397 phylogenetic HOGS. After the filtering step, 1,193 HOGs were selected to construct the superalignment matrix. Before trimming, the matrix consisted of 1,018,881 alignment positions, while after filtering it contained 473,254 (46.4%) positions which were used for the phylogenomic analysis. JTT + I + G4 + F was identified as the best-fit model and used for the phylogenetic tree inference (Figure 2). At the resulting maximum-likelihood phylogenetic tree, almost all branches were supported with 100 non-parametric bootstraps. Based on the constructed phylogeny, *P. miles* is placed within the Perciformes clade.

**Figure 2.**
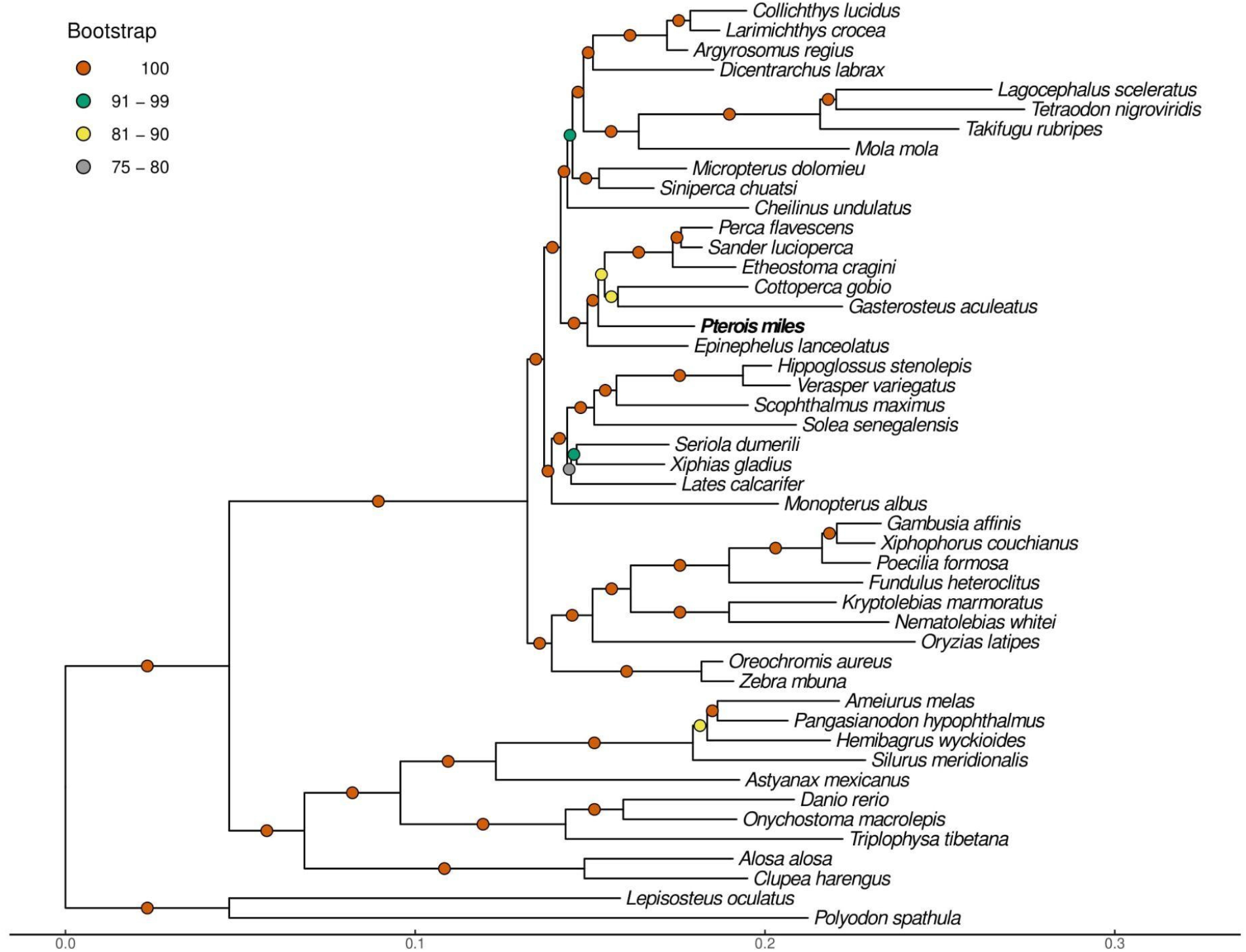
Maximum-likelihood phylogenetic tree using JTT + I + G4 + F substitution model and *P. spathula* - *L. oculatus* clade as an outgroup.

### Synteny analysis

Synteny analysis on gene level unveiled high conserved syntenic coding regions between the 42 longest contigs of *P. miles* and the 21 chromosomes of *G. aculeatus*, sharing at total 8,035 one-to-one orthologous genes (Figure 3).

**Figure 3.**
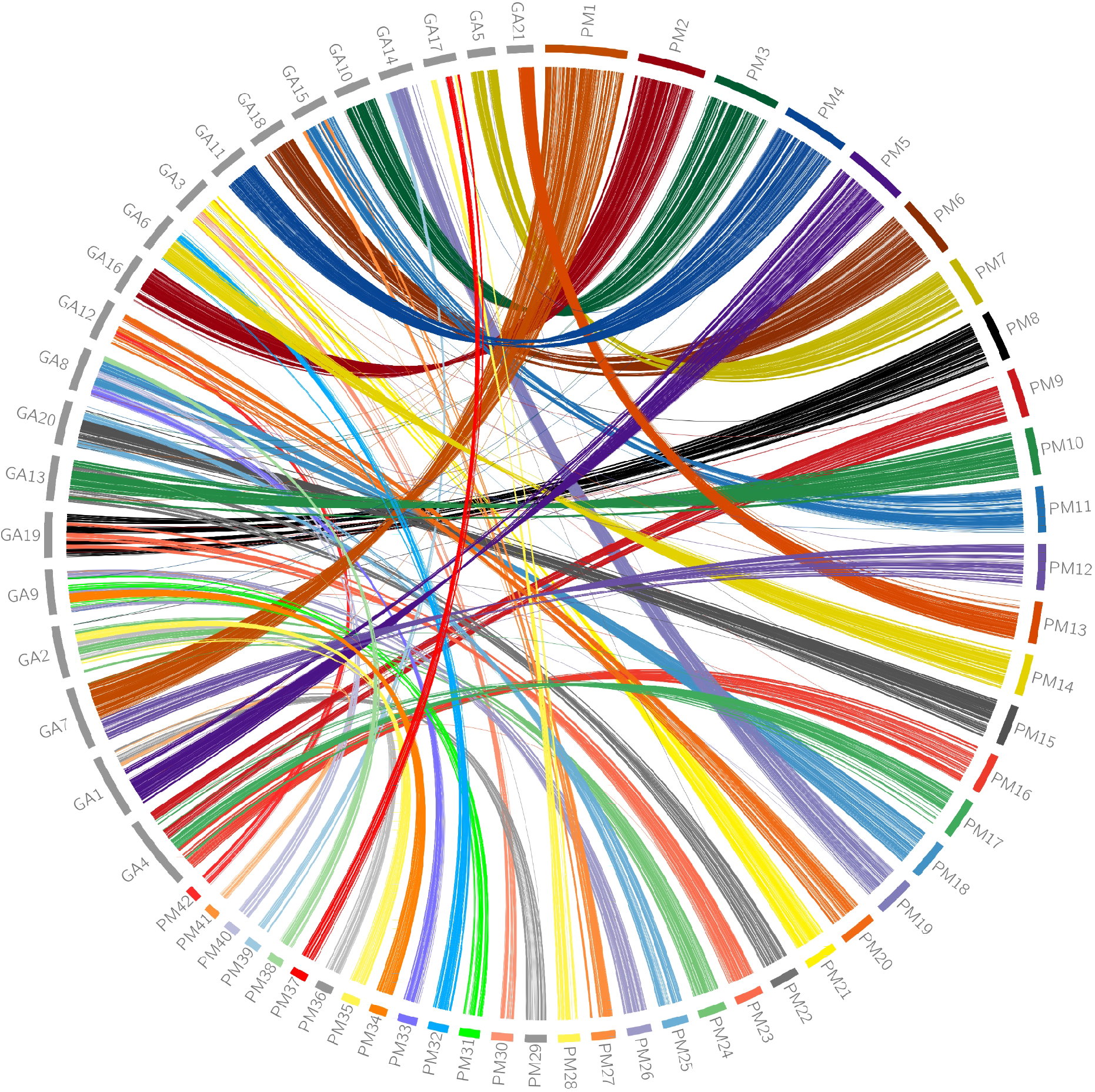
Circos plot which presents the syntenic locations of orthologous genes between the 42 longest contigs of *P. miles* (right, PM) and the 21 chromosomes of *G. aculeatus* (left, GA). Strips link orthologous genes between the two species, and colors represent the different contigs of *P. miles*.

### Gene repertoire evolution

Gene family evolution analysis in *P. miles* estimated 228 rapidly evolving gene families (out of 15,405 included families), at a significance level of 0.01 (p-value). From these rapidly evolving families, 136 were identified as expanding and 92 as contracting. The total number of genes included in these rapidly expanding families was 373. The corresponding state of the number of estimated gene families’ gains-losses inside the Perciformes species is presented in Figure 4, as a subset of Figure 2.

**Figure 4.**
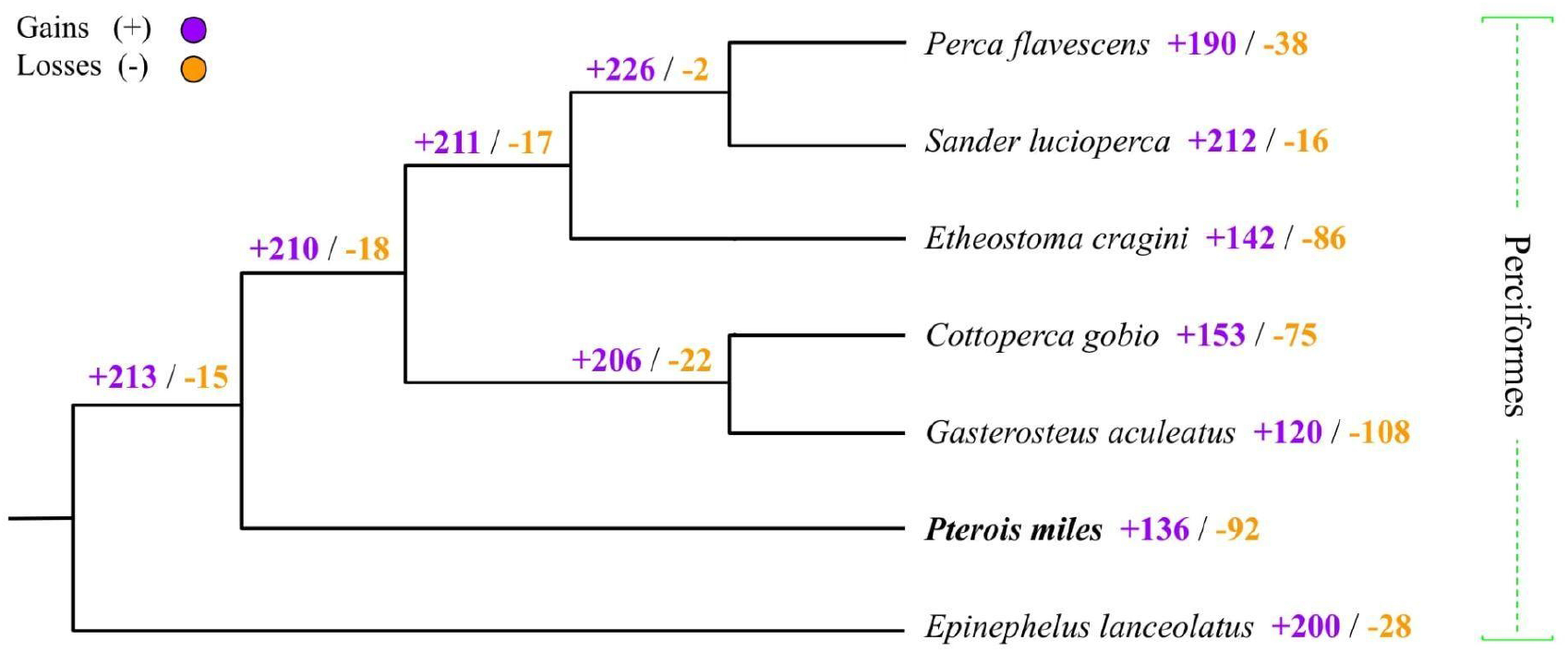
Gene family evolution analysis, including the number of gained (purple) and lost gene families (orange) for the Perciformes clade.

The number of families included in the duplication events estimation analysis was 23,775 with an average of 43 genes per family. The largest family included 1,638 genes. The total number of genes included in the gene duplication events estimation was 1,036,460. In *P. miles*, 728 gene families were identified with duplication events, including an amount of 2,263 individual genes.

The descriptive analyses for rapidly expanding gene families of CAFE and those involved in duplication events in GeneRax are presented in Figure 5. GO terms were classified into eight categories, associated with metabolism, immune system, development, growth, antimicrobial response, toxin transport, reproduction and locomotion, and the numbers of gene families, involved genes and unique terms were calculated for both results from CAFE and GeneRax. The top terms in both analyses were included in gene families associated with “metabolism”, “development”, “immune” and “growth”.

**Figure 5.**
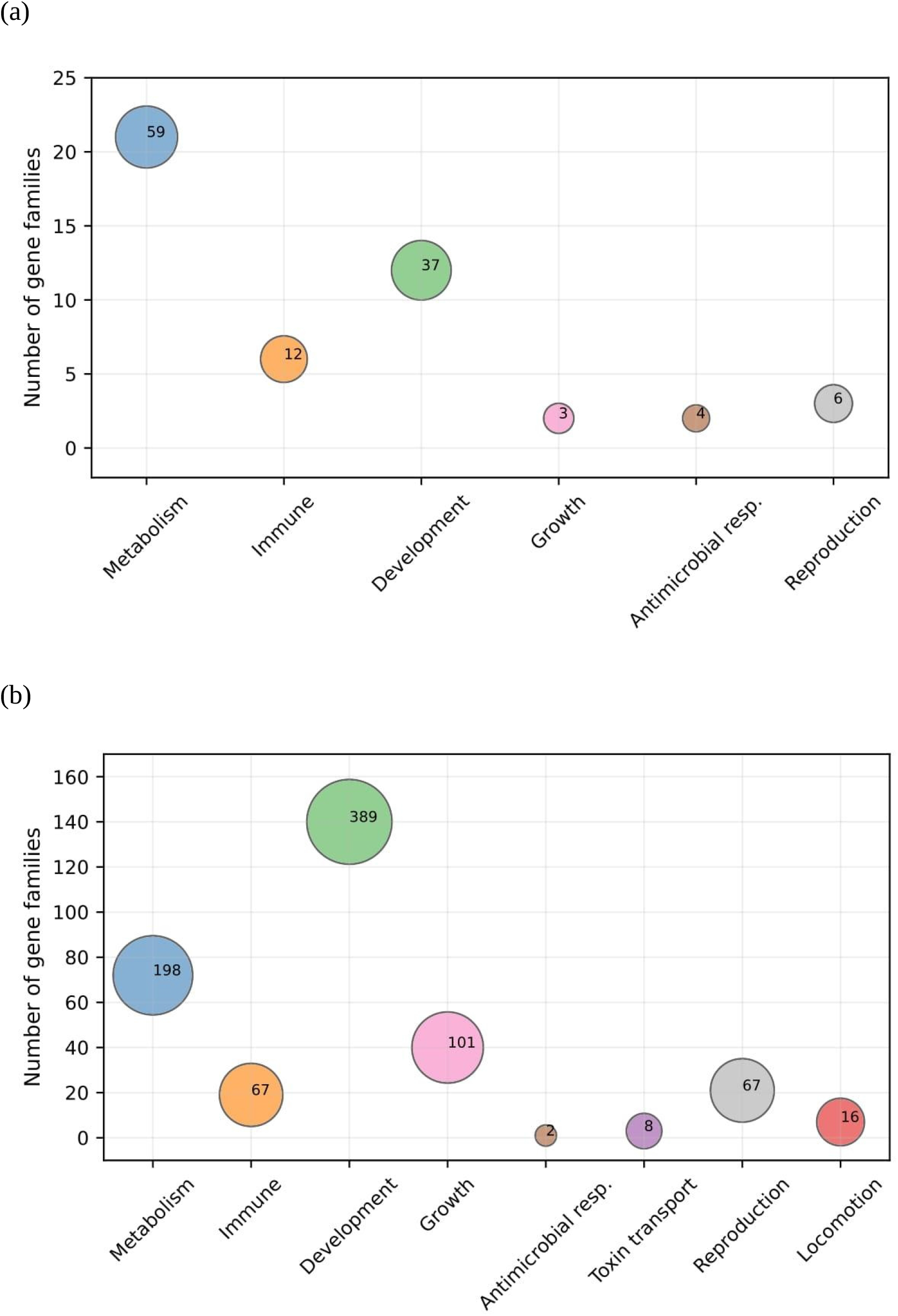
Number of orthogroups associated with specific biological processes for (a) rapidly expanding from CAFE, and (b) with duplications from GeneRax. The size of each circle is defined by the binary logarithm (log_2_) of the number of genes multiplied by the number of unique terms and then adding a scalar factor. The visualization of figures was performed with a custom python script “GO_plots.py”.

### Lionfish toxins evolution

The alignment of scorpaenid toxins protein set (blast reciprocal hits) against the genome of *P. miles* revealed a total number of six complete toxin genes, with three exons and two introns each (intron size in bp for 1: 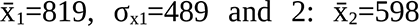 and σ_x2_=367 respectively), on the 7th longest contig and in a distance between of 50.3 kb (Supplementary_file3_pmiles_toxins.tsv). The phylogeny of scorpaenid toxins showed the separation (with high support) between the two subunits, α and β (Figure 6), which form the functional heterodimer. Also, it confirmed that three genes in devil firefish’s genome correspond to α subunit and three to β, respectively.

**Figure 6.**
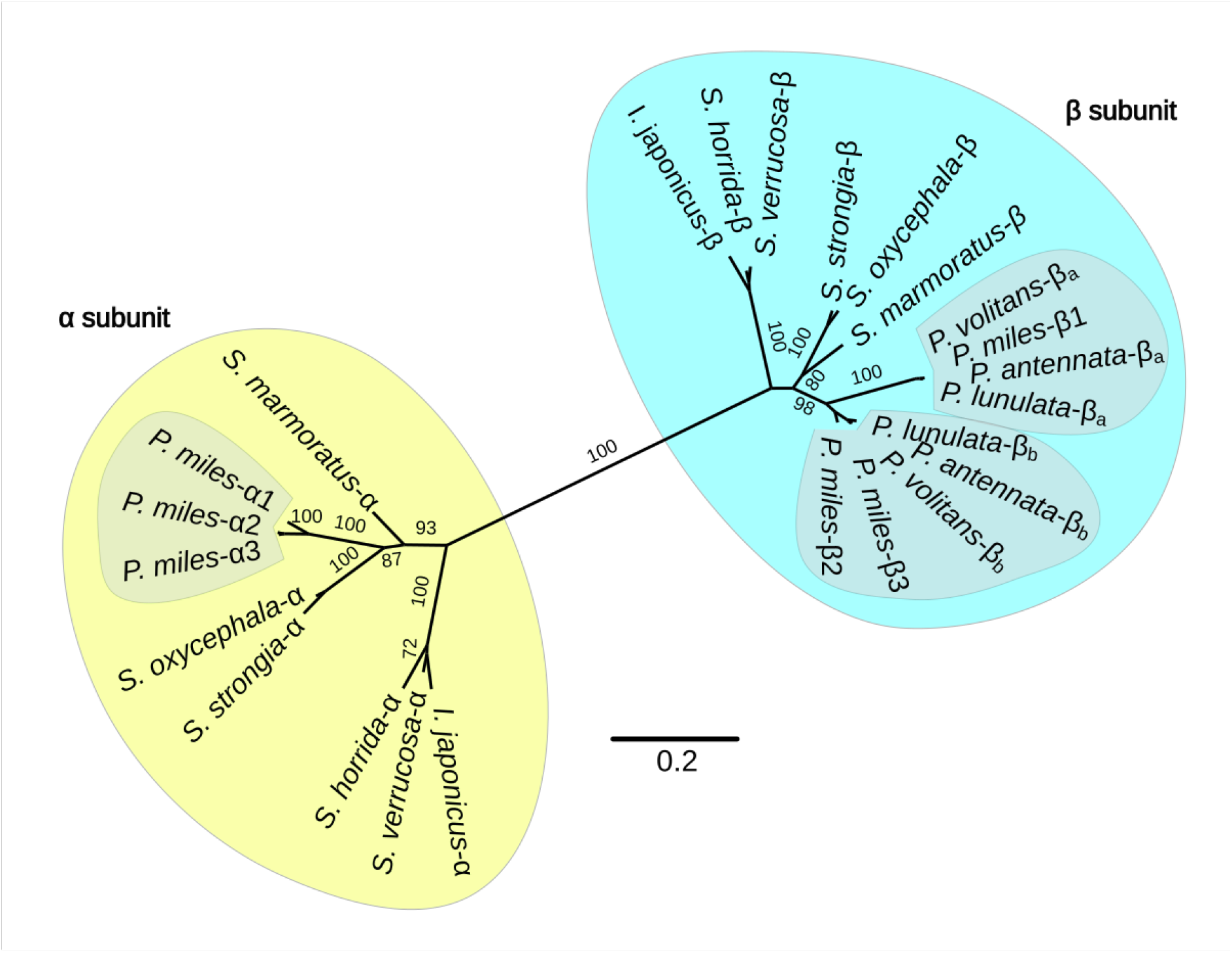
Maximum-likelihood unrooted phylogenetic tree of the two subunits of scorpaenid toxins. α subunits are presented inside the yellow and β in light blue bubble. For the phylogeny we used the JTT-DCMUT + I + G4 substitution model and conducted 100 non- parametric bootstrap replicates.

## Discussion

Here, we presented and analyzed the first high-quality genome assembly for the lessepsian migrant species *P. miles*, which is, also, the first assembly for the whole Scorpaenidae family. We positioned the species in the teleost tree for the first time and studied its complex repeat and gene content. In this study, a lionfish near-chromosome genome assembly of high-quality and contiguity was constructed from genomic data derived from three MinION flow cells and an Illumina Hiseq4000 platform, finally ending in a total size of 902.3 Mb and distributed in 660 contigs. To our knowledge, this is the first reference genome in Pteroinae and so far the only available genome of the Scorpaenidae family.

### Repeat content, gene prediction & functional annotation

The representation of TEs in the *P. miles* genome (46.5% of genome assembly) is notably higher than in other species inside Perciformes (Table 9), such as *G. aculeatus*, 13.02% (Shao et al., 2019), *S. lucioperca*, 39.0% (Nguinkal et al., 2019) and *E. lanceolatus*, 45.1% (Wang et al., 2019). Furthermore, its repetitive content is higher compared to species of similar genome size (0.9-1 Gb) such as *O. niloticus* (21.34%), *A. mexicanus* (25.21%), *O. latipes* (26.74%), *C. idella* (40.08%) and *L. oculatus* (16.06%), as presented by (Shao et al., 2019, and references therein). All aforementioned TE analyses in the different genomes, including ours presented herein, have been implemented using a different strategy for identifying the TEs of each species, a fact that could bias the results not allowing us to do a direct comparison. However, *P. miles* TE content exceeds remarkably most other fish genomes, with potential biological role in the species evolution. Elevated genome-wide repeat content has been previously linked to adaptation (Yuan et al., 2018) and invasiveness (Stapley et al., 2015; Danis et al., 2021) Thus, this higher proportion of TEs in the genome of *P. miles* could potentially play a key factor in adaptive evolution of species and consequently success in thriving in new environments. On the composition of TEs, the percentage of DNA transposons (Class II) in the assembled genome (28.63-33.8%) is only comparable to the corresponding ones in *D. rerio* (46.27% with ∼1.3 Gb genome size, Shao et al. (2019)) and *C. idella* (25.57% with ∼900 Mb genome size, Shao et al., 2019). Despite the positive correlation between genome size and the abundance of TEs in fish genomes, also confirmed here, it would be extremely interesting to investigate further the relationship between the TE heterogeneity, in terms of copy number and composition, and genomes’ evolution (Sotero-Caio et al., 2017; Shao et al., 2019). For example, 20 distinct TE superfamilies were recognised in the genome of *P. miles*, from a minimum of 77 Jockey elements to about 311,700 CACTA (Table 2), taking advantage both of the thorough classification resulting from the designed pipeline and the detailed annotation from “RM parser.py” (Supplementary Figure 1). Additionally, studies have revealed the multi-functional role of TEs on the evolution of vertebrate genomes, from genomic architecture (Sotero-Caio et al., 2017) to their relationship with non-coding RNAs (Bourque et al., 2018) and confluence to transcription regulation (Drongitis et al., 2019; Fueyo et al., 2022). Albeit, it would be noteworthy, in the superfamily level, to explore the patterns in the accumulation of TEs, their roles and consequently their contribution to gene duplication events, and genome dynamics in general.

**Table 9.** Fish genome size and TE content comparison.

| Species | Genome size (Mb) | TE content (%) | Reference |
| --- | --- | --- | --- |
| *Pterois miles* | 902.5 | 46.51 | present study |
| *Gasterosteus aculeatus* | 461.5 | 13.02 | Shao et al., 2019 |
| *Sander lucioperca* | 1014 | 39.0 | Nguinkal et al., 2019 |
| *Epinephelus lanceolatus* | 1128 | 45.1 | Wang et al., 2019 |
| *Oreochromis niloticus* | 927.3 | 21.34 | Shao et al., 2019 |
| *Astyanax mexicanus* | 1191.2 | 25.21 | Shao et al., 2019 |
| *Oryzias latipes* | 868.9 | 26.74 | Shao et al., 2019 |
| *Ctenopharyngodon idella* | 900.5 | 40.08 | Shao et al., 2019 |
| *Lepisosteus oculatus* | 945.8 | 16.06 | Shao et al., 2019 |

### Phylogenomic analysis, synteny and gene repertoire

Scorpaenidae (order: Perciformes) is a taxonomically widespread family which includes by now 370 marine species (Smith et al., 2018, and references therein), known to be venomous. Despite their worldwide distribution and diversity, this group’s biology is clearly understudied, as well as their unexplored phylogeny. Here, we presented the first phylogenetic tree that includes a representative of this family, devil firefish *P. miles*, based on whole genome data (Figure 2). This effort could be an origin for further genomic and evolutionary studies inside this family.

One-to-one orthologous genes between *P. miles* and *G. aculeatu*s, exhibited high conserved synteny (Figure 3), which confirms the high quality and completeness of constructed genome assembly. Indeed, between contigs 2, 3, 4, 6 and 7 of devil firefish and chromosomes 16, 10, 11, 18 and 5 of three-spined stickleback, there was high pairwise conservation, respectively. Taking into consideration that the haploid number of *P. miles* should be the same as its con- generic species, *P. volitans*, n=24 (Nirchio et al., 2014), an interesting fact arose. This revealed the fusion of coding regions from more than one contigs of *P. miles* to single chromosomes of *G. aculeatus* (e.g. contigs 1 and 12 to chromosome 7, contigs 5, 29, 41 to chromosome 1, contigs 9, 16, 17 to chromosome 4) and their later rearrangements (e.g. contigs 8, 23 to chromosome 19), an additional support of the high accuracy constructed assembly.

Duplication events estimation and descriptive functional analysis unveiled the extended presence of gene families, being involved in major biological processes, such as metabolism, somatic growth, immunity and reproduction (Figure 5). These families may potentially contribute to species morphology, anti-predatory tactics, rapid spread and adaptation in new marine habitats. Noteworthy, a sufficient number of immune-related gene families were identified, including immunoglobulins (Ig heavy-chain variable, light-chain variable genes), interleukins (interleukin 10 receptor), lysozymes (antimicrobial response), genes contributing to the regulation of antiviral innate immunity (e.g. TRIM35) and transcription factors that regulate the expression of MHC class II genes. An interesting finding was a detected duplication in the gene family of meprins (meprin-F in fish, Marın (2015)), proteins that are involved in toxins transport. Based on the results, it could be worth additional studies on genes responsible for the unique morphology (e.g. spines development) of devil firefish and its successful adaptation to new habitats, with the contribution of more genomic data inside the family of Scorpaenidae, that would become available in the future.

### Lionfish toxins evolution

Scorpaeniform fish toxins are multifunctional proteins that have, among others, lethal, cytolytic, hemolytic, inflammatory, nociceptive and neuromuscular activities (Campos et al., 2021). Scorpaeniform fish use their venom (toxins) mostly for defense, when the threat touches their spines (Diaz, 2015; Campos et al., 2021). These toxins are formed by two subunits α and β (Kiriake and Shiomi, 2011; Kiriake et al., 2013; Chuang and Shiao, 2014; Campos et al., 2021), being actively organized in either heterodimeric or tetrameric proteins (Campos et al., 2021). In spite of the identification of toxins in other lionfishes (*P. lunulata*, *P. volitans* and *P. antennata*), through cDNA cloning and immunoblotting (Kiriake and Shiomi, 2011; Kiriake et al., 2013), the lack of genomic data inside this genus makes it difficult to understand their relationship with other scorpaenid fish cytolysins and their evolution. We investigated the presence and identification of toxin genes on the genome, and reconstructed the phylogeny of lionfish toxins (Figure 6). The identification of three toxin genes per subunit in devil firefish and their phylogeny inside toxins from various scorpaenid fishes, rejected a previous hypothesis about the evolution of lionfish toxins. This theory proposed the absence of α subunit gene in species of genus *Pterois* and the origination of toxin genes from the β subunit of scorpaenids and a later duplication event occurred prior to the speciation of Pteroinae (Chuang and Shiao, 2014).

## Conclusion

In this study, we provide the first near-chromosome and high-quality genome assembly of devil firefish, its complex repeat and gene content, we construct the first phylogeny including a member of genus *Pterois*, based on whole genome sequencing data, and baseline the evolution of lionfish toxins. All the analyses performed here, highlighted the importance of *P. miles* genome as a valuable resource for further studies regarding the influence of transposable elements on genome evolution, the correlation between gene duplications and adaptation to new niches, lionfish rapid spread worldwide and its dominance, and scorpaenid toxins evolution.

## Supporting information

structural annotation file

functional annotation file

lionfish toxin genes structural annotation file

## Code Availability

All customs scripts, designed workflows and used software commands that have been used during this study are available at the following GitHub repositories:

- https://github.com/ckitsoulis/Pterois-miles-Genome
- https://github.com/ckitsoulis/ELDAR
- https://github.com/genomenerds/SnakeCube

## Data Availability

Genomic data from Illumina and Oxford Nanopore Technologies, and transcriptomic data from Pacific Biosciences can be accessed in the European Nucleotide Archive (ENA) under the IDS *ERR10286272*, *ERR10286273* and *ERR10286274-81* respectively. The genome assembly along with the raw data have been deposited to ENA under the study with accession PRJEB56286. Gene and functional annotations can be found in the supplementary data files.

## Acknowledgements

The authors would like to thank the scuba diver who kindly provided the specimen and the aquarists of the Cretaquarium (C.K. Doxa, T. Doulamis, P. Grigoriou, G. Vardanis, E. Pinakis) for keeping the species alive as well as for their assistance at the sampling. This research was supported through computational resources provided by IMBBC (Institute of Marine Biology, Biotechnology and Aquaculture) of the HCMR (Hellenic Centre for Marine Research)–Zorbas HPC infrastructure. Funding for establishing the IMBBC HPC has been received by the MARBIGEN (EU Regpot) project, LifeWatchGreece RI, and the CMBR (Centre for the study and sustainable exploitation of Marine Biological Resources).

