## supplementary figures and tables for "Near-chromosome level genome assembly of devil firefish, *Pterois miles*"

### SUPPLEMENTARY MATERIAL

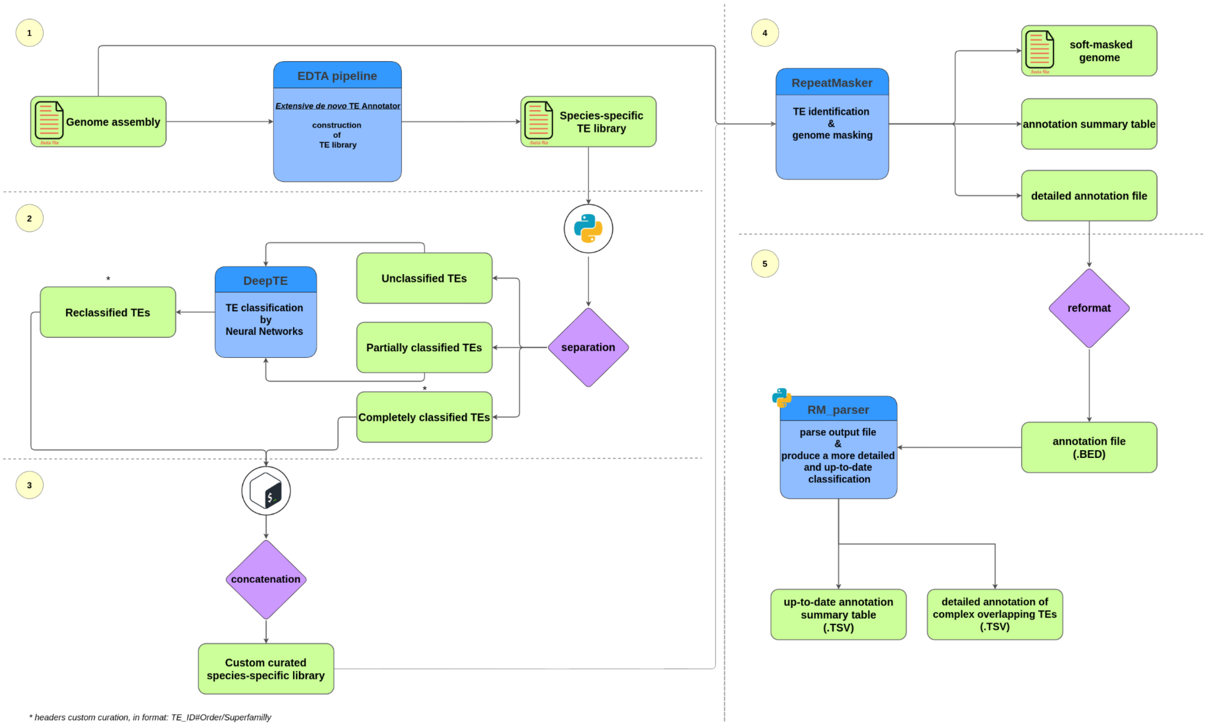


**Supplementary Figure 1.** Transposable elements annotation workflow.

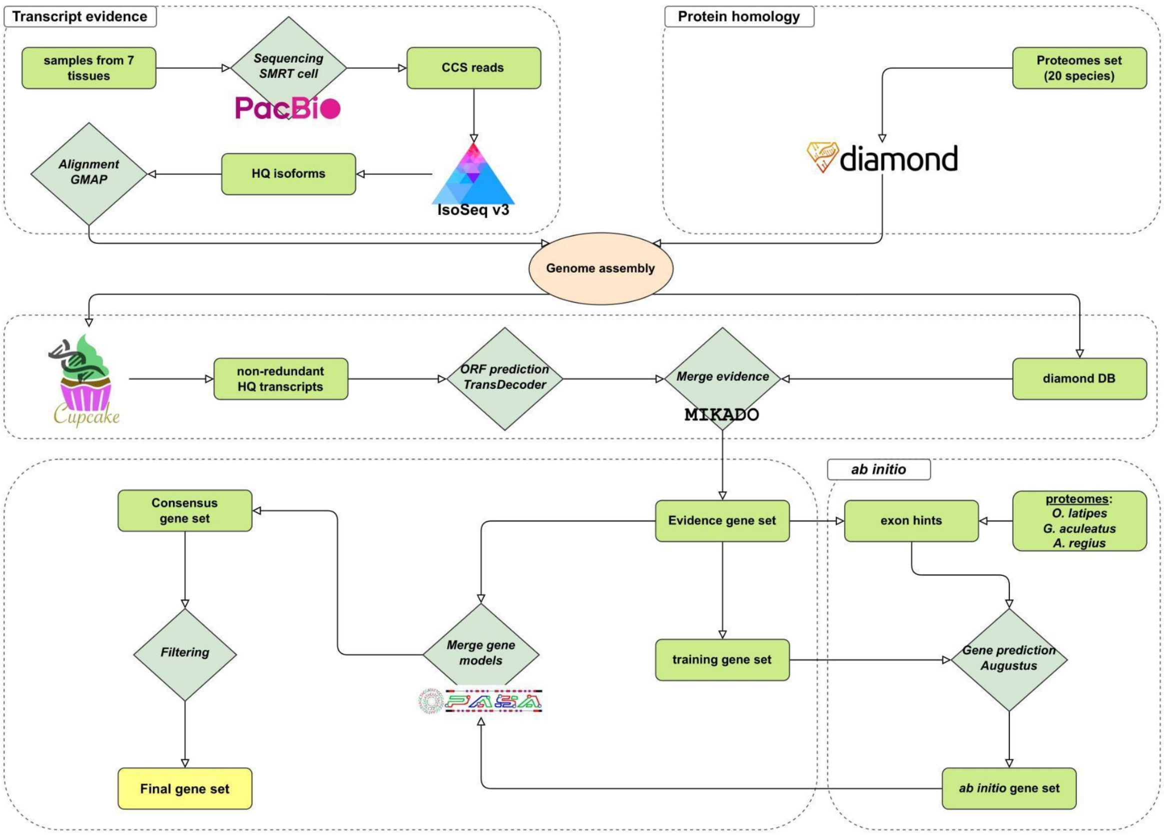


**Supplementary Figure 2.** Structural annotation workflow.

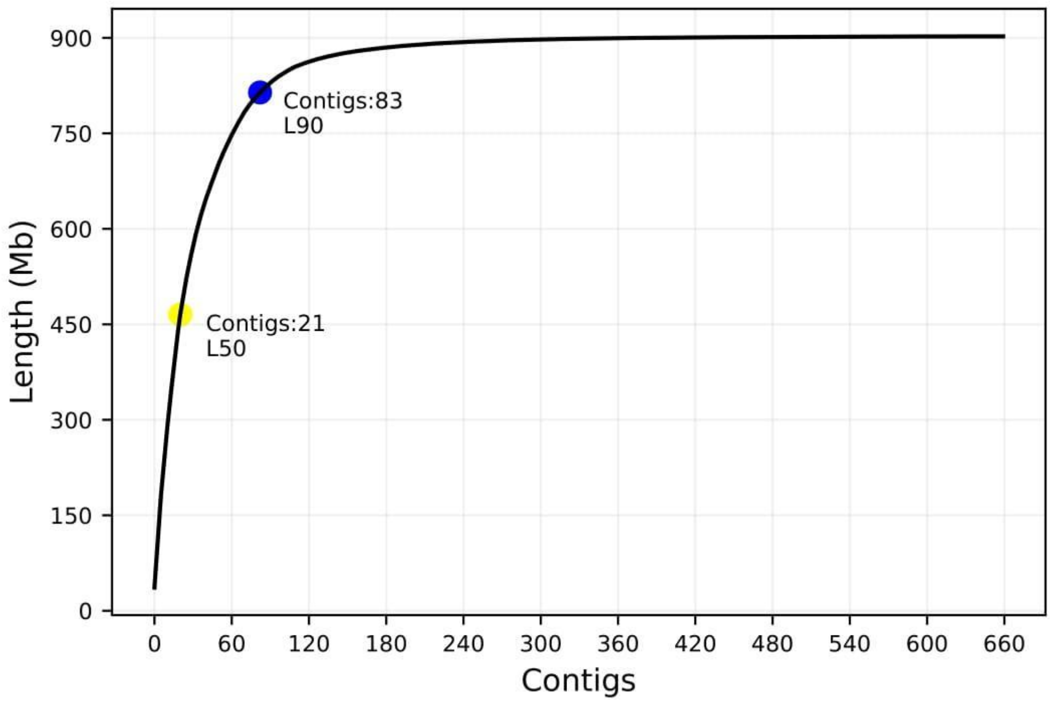


**Supplementary Figure 3.** Cumulative sum of contigs' lengths of *P. miles* genome assembly. The yellow dot shows the number of contigs that represent at least 50% (L50) and the blue one at least 90% (L90) of the genome size, respectively.

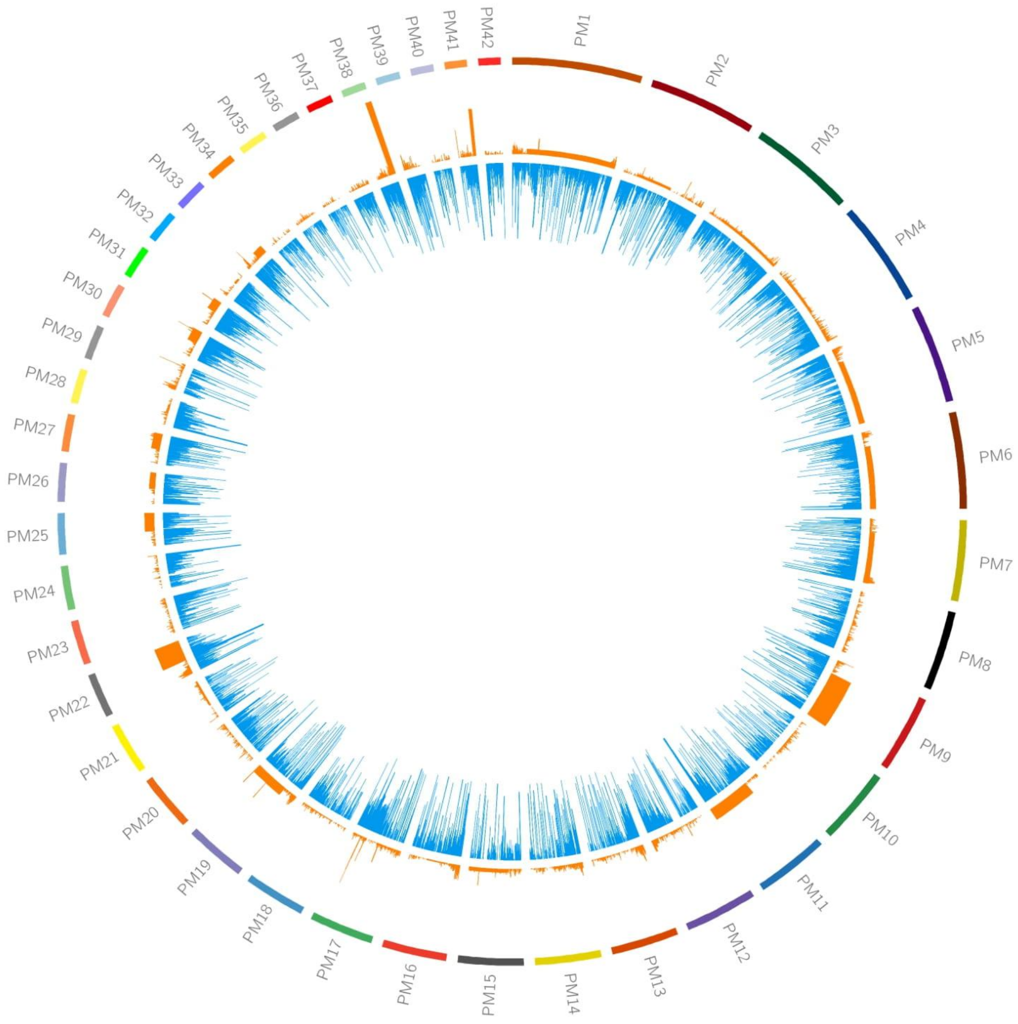


**Supplementary Figure 4.** Circos plot which presents the GC (orange - outer circle) and repetitive elements (blue - inner circle) content as histograms in the 42 longest contigs of *P. miles*, using 50kb sliding window.

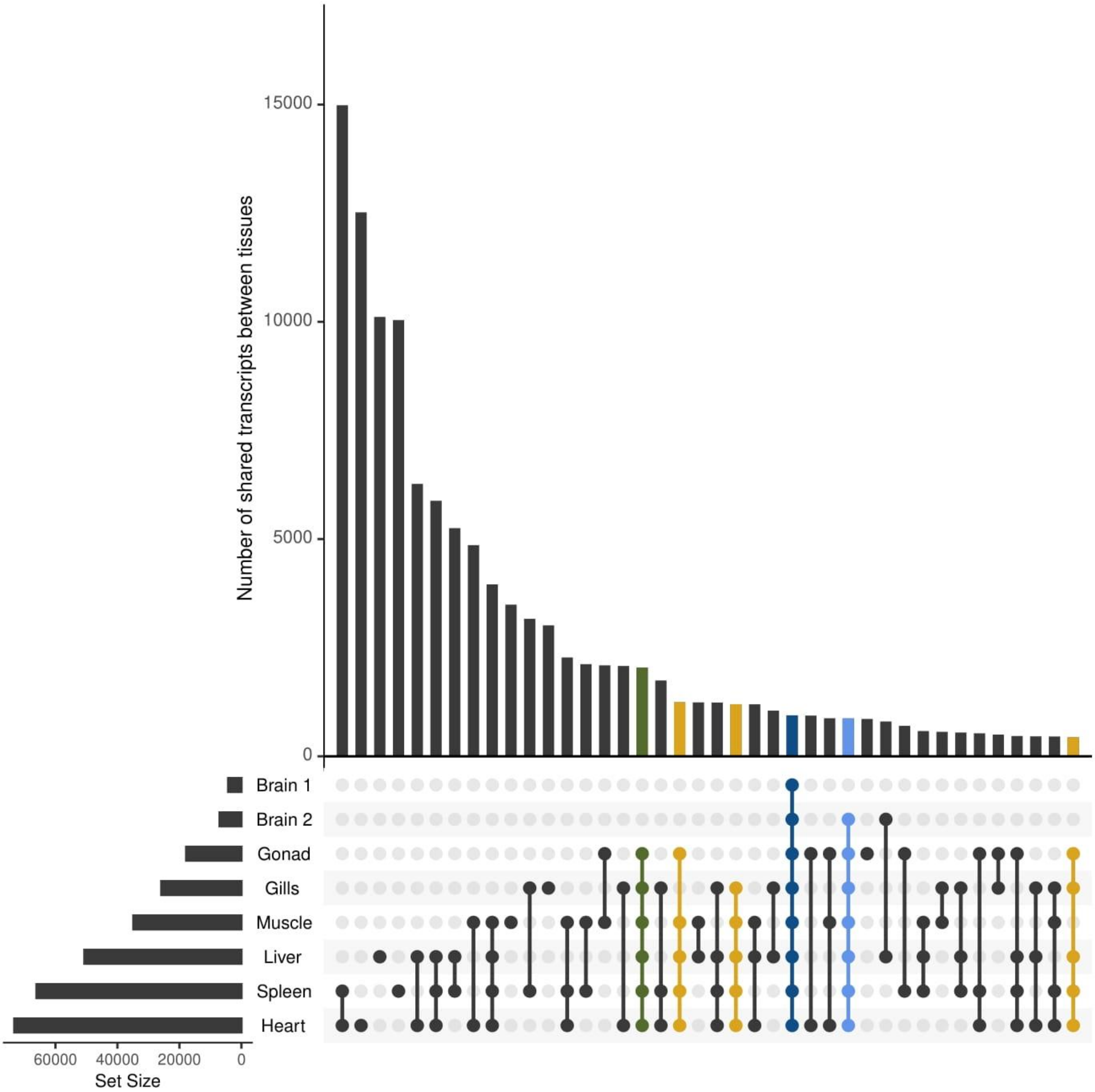


**Supplementary Figure 5.** Number of detected transcripts in each tissue and their intersections with the others using UpSetR. Blue color shows the intersection between all tissues, light blue for 7, green for 6 and yellow-gold for 5.

**Supplementary Table 1.** Species included in the phylogenomic analysis.

| Scientific name | Source | Reference | no. of proteins | BUSCO (%) |
|---|---|---|---|---|
| *Alosa alosa* | NCBI ftp | unpublished yet | 26,440 | 90.9 |
| *Ameiurus melas* | NCBI ftp | unpublished yet | 24,354 | 94.1 |
| *Argyrosomus regius* | in-house | Papadogiannis et al. (2023) | 24,443 | 95.3 |
| *Astyanax mexicanus* | Ensembl DB | McGaugh et al. (2014) | 22,998 | 97.3 |
| *Cheilinus undulatus* | NCBI ftp | Liu et al. (2021) | 23,369 | 99.2 |
| *Clupea harengus* | NCBI ftp | Kongsstovu et al. (2019) | 26,846 | 98.6 |
| *Collichthys lucidus* | NCBI ftp | Cai et al. (2019) | 28,508 | 91.7 |
| *Cottoperca gobio* | NCBI ftp | Bista et al. (2020) | 21,322 | 96.2 |
| *Danio rerio* | Ensembl DB | Howe et al. (2013) | 25,644 | 96.7 |
| *Dicentrarchus labrax* | NCBI ftp | Tine et al. (2014) | 23,380 | 95.9 |
| *Epinephelus lanceolatus* | NCBI ftp | Wang et al. (2019) | 24,223 | 99.9 |
| *Etheostoma cragini* | NCBI ftp | Reid et al. (2020) | 21,874 | 97.6 |
| *Fundulus heteroclitus* | NCBI ftp | Reid et al. (2017) | 27,033 | 99.4 |
| *Gambusia affinis* | NCBI ftp | Shao et al. (2020) | 23,272 | 99.3 |
| *Gasterosteus aculeatus* | NCBI ftp | Nath et al. (2021) | 20,779 | 98.6 |
| *Hemibagrus wyckioides* | NCBI ftp | Shao et al. (2021) | 22,794 | 95.6 |
| *Hippoglossus stenolepis* | NCBI ftp | Jasonowicz et al. (2022) | 21,840 | 98.8 |
| *Kryptolebias marmoratus* | NCBI ftp | Kelley et al. (2016) | 22,228 | 99.5 |

| | | | | |
|---|---|---|---|---|
| *Lagocephalus sceleratus* | in-house | Danis et al. (2021) | 21,333 | 91.7 |
| *Larimichthys crocea* | NCBI ftp | Ao et al. (2015) | 28,009 | 97.7 |
| *Lates calcarifer* | NCBI ftp | Vij et al. (2016) | 22,221 | 96.2 |
| *Lepisosteus oculatus* | Ensembl DB | Braasch et al. (2016) | 18,339 | 95.8 |
| *Micropterus dolomieu* | NCBI ftp | unpublished yet | 24,828 | 99 |
| *Mola mola* | NCBI ftp | Pan et al. (2016) | 21,404 | 94.1 |
| *Monopterus albus* | NCBI ftp | Tian et al. (2020) | 22,143 | 97 |
| *Nematolebias whitei* | NCBI ftp | Thompson et al. (2022) | 21,342 | 95.8 |
| *Onychostoma macrolepis* | NCBI ftp | Sun et al. (2020) | 24,754 | 93 |
| *Oreochromis aureus* | NCBI ftp | Bian et al. (2019) | 27,995 | 99.7 |
| *Oryzias latipes* | NCBI ftp | Kasahara et al. (2007) | 23,620 | 96.4 |
| *Pangasianodon hypophthalmus* | NCBI ftp | Gao et al. (2021) | 21,245 | 93.5 |
| *Perca flavescens* | NCBI ftp | Feron et al. (2020) | 23,990 | 99.6 |
| *Poecilia formosa* | NCBI ftp | Warren et al. (2018) | 23,165 | 98.6 |
| *Polyodon spathula* | NCBI ftp | Cheng et al. (2020) | 30,763 | 97.2 |
| *Pterois miles* | in-house | current study | 24,639 | 93.8 |
| *Sander lucioperca* | NCBI ftp | Nguinkal et al. (2019) | 25,044 | 99.7 |
| *Scophthalmus maximus* | NCBI ftp | Xu et al. (2020) | 21,737 | 99.5 |
| *Seriola dumerili* | NCBI ftp | Araki et al. (2018) | 23,276 | 98 |
| *Silurus meridionalis* | NCBI ftp | Zheng et al. (2021) | 22,769 | 95.2 |

| | | | | |
|---|---|---|---|---|
| *Siniperca chuatsi* | NCBI ftp | Ding et al. (2021) | 22,756 | 99 |
| *Solea senegalensis* | NCBI ftp | Guerrero-Cózar et al. (2021) | 23,462 | 99.3 |
| *Takifugu rubripes* | Ensembl DB | Aparicio et al. (2002) | 21,411 | 93.8 |
| *Tetraodon nigroviridis* | Ensembl DB | Jaillon et al. (2004) | 19,600 | 88.6 |
| *Triplophysa tibetana* | NCBI ftp | Yang et al. (2019) | 24,310 | 93.1 |
| *Verasper variegatus* | Ensembl DB | Zhao et al. (2021) | 21,273 | 97.8 |
| *Xiphias gladius* | NCBI ftp | Wu et al. (2021) | 21,527 | 99.7 |
| *Xiphophorus couchianus* | NCBI ftp | Shen et al. (2016) | 22,879 | 99.7 |
| *Zebra mbuna* | NCBI ftp | Conte and Kocher (2015) | 26,063 | 99.6 |
